# Deconstructing the structural conservation of distantly related bacterial nucleoid-associated proteins using functional chimeras

**DOI:** 10.1101/804138

**Authors:** Rogério F. Lourenço, Saumya Saurabh, Jonathan Herrmann, Soichi Wakatsuki, Lucy Shapiro

**Affiliations:** Department of Developmental Biology, Stanford University School of Medicine, Stanford, California; Department of Structural Biology, Stanford University School of Medicine, Stanford, California; Bioscience Division, SLAC National Accelerator Laboratory, Menlo Park, California; Chan Zuckerberg Biohub, San Francisco, California

**Keywords:** nucleoid-associated protein, AT-rich DNA loci, chimeras, structure/function conservation, bridging DNA-binding mode

## Abstract

Nucleoid-associated proteins (NAPs) are DNA-binding proteins critical for the organization and function of the bacterial chromosome. A subclass of NAPs, including *Caulobacter crescentus* GapR and *Escherichia coli* H-NS, preferentially bind AT-rich regions of the nucleoid, but phylogenetic groups that encode GapR rarely encode H-NS. Here, utilizing genetic, biochemical, and biophysical studies of GapR in light of a recent DNA-bound crystal structure of GapR (Guo et al, 2018), we show that although evolutionarily distant, GapR and H-NS possess two regions that are structurally and functionally conserved. These regions are involved in self-association and DNA-binding, even though the two proteins oligomerize and regulate transcription differently. Functional analysis of GapR and H-NS protein chimeras identified structural elements present in H-NS but absent in GapR that rationalize differences in transcriptional regulation. In addition, we identified a sequence element unique to GapR that enables assembly into its tetrameric state. Using fluid-atomic force microscopy, we showed that GapR is capable of bridging DNA molecules *in vitro*. Together, these results demonstrate that two distantly related NAPs utilize evolutionarily conserved structural elements to serve specialized cellular roles via distinct mechanisms.

## INTRODUCTION

Bacterial cells compact and organize their genetic material in a hierarchically ordered and dynamic structure called the nucleoid (1). This complex organization of DNA is achieved in part by the action of an array of nucleoid-associated proteins (NAPs) that affect multiple cellular processes including replication, transcription, and recombination (2). Except for an intrinsic high-affinity for DNA, NAPs are notably diverse with respect to their structures, the mechanisms by which they recognize DNA, and the extent to which they affect DNA-related processes (2). The histone-like nucleoid structuring protein H-NS is remarkable due to its ability to silence AT-rich sequences (3–7). By spreading across promoter sequences, H-NS is thought to prevent RNA polymerase from accessing these regions (5). Further, H-NS filaments assemble at sites of Rho-dependent termination, where they bridge segments of the nucleoid and contribute to both the pausing and termination of transcription (8).

Members of the H-NS family have been widely recognized in the β and γ subdivisions of Proteobacteria (9). Even though some NAPs in distantly related bacteria, such as Lsr2 in *Mycobacterium tuberculosis* and the Rok protein in *Bacillus subtilis*, display low sequence similarity to H-NS, they exhibit the same preference for DNA with high AT content and can functionally replace H-NS in *Escherichia coli* (10–12). In addition, Lsr2 bridges DNA as does H-NS (13). Regardless of whether Lsr2 and the Rok protein are H-NS orthologs or represent convergent evolution, NAPs that specifically recognize AT-rich sequences are more widely spread in bacteria than can be assumed by only considering proteins with clear sequence similarity to H-NS.

GapR, a newly discovered NAP with binding preference for regions with high AT content, was identified in the asymmetrically dividing bacterium *Caulobacter crescentus* and was found to have a broad distribution in the α subdivision of Proteobacteria (14–17). Interestingly, over-occupation of high AT chromosome sites in *C. crescentus* due to high expression of either GapR or H-NS from *E coli*, causes morphological and division defects (16). However, association of GapR with these sites does not lead to downregulation of gene expression (16). Furthermore, *gapR* expressed in *E. coli* fails to restore the ability of an *h-ns* mutant to repress expression of the *bgl* operon (16). Therefore, although both GapR and H-NS occupy AT-rich sequences, they diverge with respect to the downstream effects on gene expression.

Recently, GapR was shown to associate with the 3’ ends of highly transcribed genes, which do not contain AT-rich sequences but are generally over-twisted due to DNA unwinding (15). A high-resolution crystal structure of *Caulobacter* GapR has revealed that it can assemble into a tetramer that encircles over-twisted DNA. DNA-binding by GapR stimulates type-II topoisomerases to relax positive supercoiling (15). Therefore, unlike H-NS, GapR does not block the access of the transcription machinery to DNA and seems to lack the structural elements required to form a higher-order oligomer.

In this study, we engineered chimeric proteins by combining parts of H-NS and the GapR protein. Using this approach, we identified two regions of functional similarity between GapR and H-NS; one of them responsible for the self-association of the proteins in a coiled-coil structure, and the other important for binding DNA. We also identified key H-NS structural elements that are missing in GapR, which may account for the observations that GapR does not form filaments in the absence of DNA nor represses transcription when expressed in *E. coli*. Notably, GapR was found to have a unique sequence element that allows the protein to assemble into a tetramer. Despite these remarkable differences, GapR, like H-NS, can bridge DNA.

## MATERIALS AND METHODS

### Bacterial strains and growth conditions

Strains used in this work are listed in Table S1. *E. coli* strains were grown in Luria broth at 37°C for all experiments with the following exceptions: i) cells were grown in MacConkey plates containing 1% maltose for analysis of *in vivo* protein-protein interaction in a bacterial two-hybrid system; ii) MacConkey plates supplemented with 0.5% salicin were used for the *bgl* complementation assay; iii) protein expression was conducted at 30°C. When appropriate, the growth medium was supplemented with 0.02% L-arabinose, 50 µg/mL ampicillin, 30 µg/mL kanamycin, 20 µg/mL chloramphenicol, 12 µg/mL tetracycline, 50 µg/mL spectinomycin and/or 30 µg/mL streptomycin. Experiments with *C. crescentus* were performed using CB15N-derivatives grown at 22°C or 30°C in peptone-yeast extract (PYE) medium supplemented or not with 25 µg/mL spectinomycin and 5 µg/mL streptomycin. Cell growth was monitored by measuring the optical density at 600 nm, and aliquots were removed, serially diluted (1:10 dilution) and plated on PYE medium for counting colony-forming units.

### Plasmid construction

Plasmids used in this work were constructed by the Gibson Assembly method (18) and are described in Table S2. Plasmids linearized with restriction enzymes were combined with PCR products generated using oligonucleotides comprising a sequence annealing to the region to be amplified and a flanking region of homology to the target vector or to another fragment when more than two DNA fragments were assembled together in the same reaction. For random mutagenesis, the *gapR* coding sequence corresponding to the amino acid residues 1-52 was amplified by error-prone PCR as previously described (19). Site-directed mutagenesis was performed using oligonucleotides modified to include the specific amino acid substitution. Plasmids were introduced into *E. coli* and *C. crescentus* by electroporation. Sequences of the oligonucleotides are available on request.

### Gene replacement in *C. crescentus*

pNPTS138-derivatives were constructed with the spectinomycin/streptomycin resistance gene and its promoter inserted upstream from the wild-type or a mutant copy of *gapR* along with its native promoter. The fragments flanking each side of the *gapR* locus were included in order to allow the replacement of WT *gapR* in CB15N with a mutant copy of the gene and the insertion of the sequence conferring resistance to spectinomycin and streptomycin by two homologous recombination events. Mutant strains were isolated by analyzing colonies with respect to the restriction profile of PCR-amplified *gapR* followed by sequencing of the entire region using oligonucleotides annealing outside the region cloned into pNPNTS138.

### Phase contrast microscopy

*C. crescentus* cells grown to exponential phase (0.3 < OD_600 nm_ < 0.5) were diluted to OD_600 nm_ = 0.1, and 1 µL was spotted on agarose pads containing M2G minimal medium. Cells were imaged by phase contrast microscopy using an inverted microscope (DMi800, Leica) equipped with a 100 x (1.4 N.A.) oil objective.

### RNA extraction and quantitative RT-PCR

*E. coli* cultures diluted to OD_600 nm_ = 0.1 were grown for 24 h, and cells were harvested (16,000 x g for 1 min). Cell pellets were suspended in 1 mL Trizol Reagent (Thermo Fisher Scientific), and total RNA was extracted according to manufacturer’s instructions. Samples were treated with DNAse I (Thermo Fisher Scientific) and their integrity was checked by agarose gel electrophoresis. 2.5 µg of DNA-free RNA samples were used as template for cDNA synthesis in the presence of 0.2 µg random hexamer primer, 1 mM dNTP mix and 200 U RevertAid Reverse Transcriptase (Thermo Fisher Scientific). Real-time PCR was performed using 0.4 µM of each gene-specific oligonucleotide and 1 X Fast SYBR Green Master Mix (Applied Biosystems). Fluorescence was monitored by the 7500 Fast Real-Time PCR System (Applied Biosystems). Oligonucleotide sequences were designed with the Primer3 software version 0.4.0 (20) and are available on request. The 2^-ΔΔCT^ method (21, 22) was utilized to calculate relative expression of genes, normalized to the *rpoD* gene (23).

### Protein expression and purification

GapR was expressed in *E coli* BL21(DE3) strain by growing cells for 3 h at 30°C in the presence of 0.5 mM isopropylthiogalatoside (IPTG). Cells were harvested, washed twice in buffer A (20 mM HEPES-NaOH pH 7.5, 150 mM NaCl, 10% glycerol), snap frozen in liquid nitrogen and stored at -80°C. For protein purification, cell pellets were thawed and resuspended in buffer A supplemented with 25 mM imidazole, 1 mg/mL lysozyme and EDTA-free protease inhibitor cocktail (Santa Cruz Biotechnology). After 1 h treatment at 4°C, cells were further disrupted by sonication. Lysates were cleared by two rounds of centrifugation at 21,000 x g, and cleared lysates were incubated with Nickel-nitrilotriacetic acid (Ni-NTA) agarose (ThermoFisher Scientific) pre-equilibrated with the same buffer used for cell lysis. The agarose beads were collected and washed three times with buffer A supplemented with increasing concentration of imidazole (25, 50, 100, 200 and 500 mM). Different GapR proteins were eluted using distinct concentrations of imidazole. Proteins were concentrated using Amicon ultra centrifugal filter MWCO 3kDa (Millipore), dialyzed against buffer B (20 mM HEPES-NaOH pH 7.5, 1 M NaCl, 1 mM EDTA, 10% glycerol) and run in the size exclusion chromatography column Superdex 200 10/300 GL (GE Healthcare Life Sciences) to separate most of the co-purified DNA. After purification, the proteins were concentrated, evaluated by SDS-PAGE, snap frozen in liquid nitrogen and stored at -80°C. The proteins were thawed and used directly or dialyzed to buffer A prior to the usage. Protein quantification was carried out using Bradford reagent (BioRad). Both analytical and preparative chromatography analyses were performed using the NGC chromatography system (Bio-Rad).

For SAXS analysis, fresh proteins purified by affinity and size-exclusion chromatography as described above were further purified by cation exchange chromatography using the HiTrap SP HP column (GE Healthcare Life Sciences). For cation exchange chromatography, the proteins were dialyzed against buffer C (20 mM HEPES-NaOH pH 7.5, 50 mM NaCl, 10% glycerol), and elution was carried out using 50% buffer D (20 mM HEPES-NaOH pH 7.5, 1 mM NaCl, 10% glycerol). The proteins were then dialyzed against buffer A, quantified and used directly for SAXS measurements.

### Microscale thermophoresis analysis

For the thermophoresis assays, a fluorescently-labeled double strand DNA was PCR-amplified or obtained by hybridization. In each case, an oligonucleotide covalently bound at its 5’ end to the ATTO-488 fluorophore was used. The hybridization was carried out by heating the oligonucleotides at 95°C for 3 min followed by incubation at 55°C for 3 min. 0.1 µM labeled DNA was mixed with protein in 20 mM HEPES-NaOH pH 7.5, 150 mM NaCl, 10% glycerol, 0.05 % Tween-20, and the resulting mixture was serially diluted (1:1 dilution) in the same buffer supplemented with fluorescently-labeled DNA. After 30 min incubation at room temperature, samples were analyzed in the Monolith NT.115 instrument (Nano Temper Technologies). The fluorescence was monitored for 5 s before heating, for 30 s under constant heating (LED Power 80% and MST Power 40) and for 5 s after deactivating the infrared laser. A 25 s delay was allowed between successive measurements. Data analysis was carried out using the software MO Affinity Analysis version 2.2.4 (Nano Temper Technologies).

### Electrophoretic mobility shift assay and native PAGE

The electrophoretic mobility of proteins under native conditions and the electrophoretic mobility shift of DNA upon incubation with proteins were determined using gel electrophoresis on 4-15% polyacrylamide gels (BioRad). Protein samples and loading buffer (20 mM HEPES-NaOH pH 7.5, 150 mM NaCl, 30% glycerol, 0.05% bromophenol blue) were mixed at a ratio of 1:1 (v/v) and run at 4°C in 1 X Tris-glycine buffer (25 mM Tris-Cl pH 8.3, 192 mM glycine). For electrophoretic mobility shift assays, PCR-amplified DNA was incubated with protein for 30 min at room temperature in 20 mM HEPES-NaOH pH 7.5, 10% glycerol supplemented with 150 or 500 mM NaCl and then subjected to electrophoresis as described above.

### Protein crosslinking

Crosslinking reactions were performed by incubating proteins with 400 mM 1-ethyl-3-(-3-dimethylaminopropyl) carbodiimide hydrochloride (EDC) and 100 mM N-hydroxysulfosuccinimide (NHS) for 2 h in 20 mM HEPES-NaOH pH 7.5, 150 mM NaCl, 10% glycerol. Crosslinking reactions were stopped by adding 150 mM β-mercaptoethanol, 0.1% SDS and heating at 95°C for 5 min. Samples were then mixed with SDS loading buffer, boiled for 5 min and resolved on 4-15% polyacrylamide gels (BioRad) under denaturing condition (25 mM Tris-Cl pH 8.3, 192 mM glycine, 0.1% SDS).

### Small angle X-ray scattering measurement

Small angle X-ray scattering (SAXS) experiments were performed at the bio-SAXS beamline BL4-2 at the Stanford Synchrotron Radiation Light source. Data were collected using a Pilatus3 X 1M detector (Dectris AG) with a 3.5 m sample-to-detector distance and beam energy of 11 keV (wavelength, *λ* = 1.127 Å). SAXS data were measured in the range of 0.0033 Å^−1^ ≤ *q* ≤ 0.27 Å^−1^ (*q* = 4*π*sin(*θ*)/*λ*, with 2*θ* being the scattering angle). The *q* scale was calibrated with silver behenate powder. The GapR samples were injected directly into a temperature-controlled flow cell. The SAXS data were taken in a series of 12, 1 s exposures. These images were then analyzed for possible effects of radiation damage, normalized according to the transmitted intensity, and averaged using the program SasTool (http://ssrl.slac.stanford.edu/~saxs/analysis/sastool.htm).

### Circular dichroism spectroscopy

Circular dichroism (CD) measurements were performed using a J-815 Circular Dichroism Spectrometer (Jasco). Far-UV spectra (200-250 nm) were recorded in a 1 mm path-length cell with an exposure time of 1 s/nm. The sample cell was maintained at 15°C and three scans were collected and averaged for each sample. A buffer spectrum was subtracted from all sample spectra before plotting.

### Atomic force microscopy

DNA molecules used for AFM experiments were composed of the *pilA* promoter (316 bp) and its downstream region (704 bp) and were obtained by amplification using PCR reactions. DNA was incubated alone or in the presence of GapR (wild-type and a mutant protein) for 16 h at 4°C and then imaged on BioScope Resolve Bio AFM (Bruker). The long incubation time was used to assure that the binding reaction reached the equilibrium. The microscope itself was placed in an isolation box to minimize drift, temperature fluctuations and vibrations. Rapid force-distance (PeakForce Tapping mode) imaging modality was used to obtain images. We used a high-resolution AFM probe (PeakForce HIRS-F-B, spring constant k = 0.12 N/m) for imaging. After deploying the probe on the AFM head, a laser alignment was performed followed by a thermal tuning calibration of the probe in MilliQ water. For high resolution AFM of DNA and DNA-GapR complexes, we used freshly cleaved mica as a substrate. Mica was cleaved by removing five to six layers using scotch tape. To minimize sample drift, we affixed cleaved mica on steel specimen discs (Ted Pella, Inc) using optical glue (NOA68, Norland optical adhesives). The optical glue also created a hydrophobic barrier around the mica, thus creating a sample well that minimized sample overflow and evaporation. The sample chamber was then mounted on the microscope using magnetic mounts. Wet wipes were kept around the chamber and AFM head to further minimize evaporation. 100 µL of imaging buffer containing 20 mM HEPES-NaOH pH 7.5, 150 mM NaCl, 10% glycerol, 10 mM NiCl_2_ was added to the sample chamber. Next, 20 µL of sample (DNA or DNA + GapR) was added to the imaging chamber. After addition of the sample, we pipetted 20 µL of the imaging solution up and down five times to ensure uniform mixing. This procedure led to a very reproducible sample density across different fields of view. After mixing the sample, the aligned AFM head was carefully placed on the microscope and the cantilever was moved down until it was submerged in the sample. At this stage another round of laser guided calibration and thermal tune calibration was performed. Finally, the AFM cantilever was engaged with the sample and we waited for 5-20 min before changing any parameters to equilibrate the system and improve stability. We scanned a 1 μm by 1 μm region to obtain multiple DNA molecules in one field of view. Nanoscope software was used for data acquisition. Typically, we acquired 15-20 images with the high-resolution tip before observing a deterioration in the data quality. Due to this limitation, we used a new tip for each sample, keeping all the scanning parameters the same. For analyzing AFM data, we developed a workflow that comprised of a first round of image processing (image flattening using a 0^th^, 1^st^ and 2^nd^ order polynomial applied to each scan line) followed by particle segmentation using sample heights as a cut-off. The cut-off was selected such that we were able to threshold > 90% of DNA molecules in the image. Image processing and segmentation were performed in Bruker NanoScope Analysis software v 9.0. Post-segmentation, we obtained heights of DNA (or DNA + GapR) segments and exported them into text files that were processed using bespoke programs written in Matlab (MathWorks).

## RESULTS

### GapR and H-NS proteins share regions that are structurally and functionally similar

Both GapR (14–16) and H-NS (3–7) are NAPs that preferentially bind to AT-rich regions of the chromosome. Comparison of the GapR and H-NS primary structures revealed two regions of the GapR sequence that display clear similarity to H-NS (Fig. 1A): 1) The N-terminal region of GapR (residues 2-49) displays 27% identity and 61% similarity to residues 4-52 of H-NS. In each case, a coiled-coil motif lies within these N-terminal regions (Fig.1A). These sequences adopt a dimeric, antiparallel coiled-coil structure referred to as dimerization site 1 in H-NS (15,24–26) and Helix 1 in GapR (15,24–26) (Fig. 1A). 2) The segment of GapR that encompasses Helix 2 of the protein (residues 50-70) (15) shares 29% identity and 67% similarity with a part of the DNA-binding domain of H-NS (residues 111-129) (27) (Fig. 1A). Moreover, this comparison also revealed two main differences between GapR and H-NS. 1) The central region of H-NS (residues 53-110), which corresponds to another dimerization site of the protein (dimerization site 2) (24, 26) and a segment of the DNA-binding domain including a flexible linker proposed to mediate non-specific DNA-association (28), is completely absent in GapR (Fig. 1A). 2) Helix 3 of GapR (residues 71-89) appears to represent a unique structural element (Fig. 1A).

**Figure 1.**
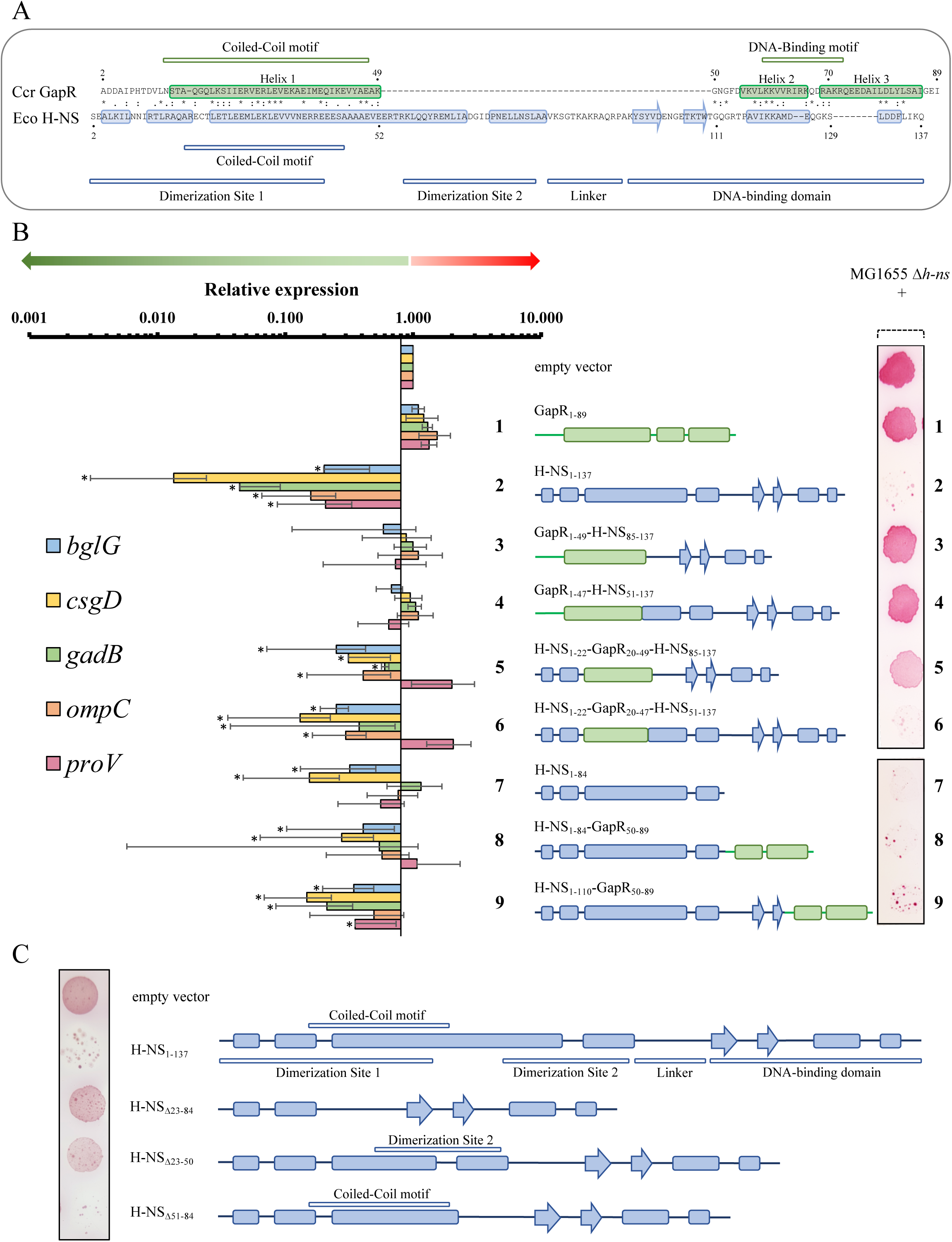
GapR and H-NS share two functionally similar regions. **A)** Comparison of the deduced amino acid sequences of *C. crescentus* GapR and *E. coli* H-NS. The sequences were compared by Clustal Omega (48), and the alignment was refined manually. Structurally determined elements are shown as boxed regions for α-helices and boxed regions with arrowheads for β-sheets (shown in green for GapR and blue for H-NS). Green bars shown above the GapR sequence correspond to the previously reported coiled-coil and DNA-binding motifs (14). Blue bars shown below the *E. coli* H-NS sequence correspond to the coiled-coil motif, the dimerization sites 1 and 2, the flexible linker and the DNA-binding domain (24–28). The N-terminal methionine was omitted from both amino acid sequences in order to allow a more precise alignment. **B)** Ability of chimeric proteins to repress gene expression. The right panel shows diagrams of the *Caulobacter* GapR (green), *E. coli* H-NS (blue) and chimera constructs, numbered 1-9, next to the ability of strains carrying these constructs to repress the *bgl* operon in Δ*h-ns E coli*. MG1655 Δ*h-ns E coli* (28) carrying derivatives of the arabinose-inducible expression vector pBAD33 were grown in the presence of 0.02% arabinose, and 3 µL from each culture were spotted on a MacConkey plate supplemented with 0.5% salicin and 0.02% arabinose, incubated at 37°C for 24 h, and imaged (49). H-NS, but not GapR, was able to repress the *bgl* operon. The left panel shows values representing the fold change in gene expression in the corresponding chimera strains compared to cells harboring the empty vector. qRT-PCR experiments were performed with total RNA extracted from cells bearing each chimera grown for 24 h in the presence of 0.02% arabinose. Results were normalized using the *rpoD* gene as an endogenous control. Data are mean values of three independent experiments; bars represent the standard deviation. Values considered statistically significant (p<0.001 using unpaired Student’s t-Test) are denotated by asterisks. Color-coded bar graphs of the expression levels of 5 genes known to be repressed by WT H-NS (*bglG, csgD, gadB, ompC and proV*) are shown for each of the chimeras. **C)** Dimerization site 1 but not dimerization site 2 is critical for the ability of H-NS to repress the *bgl* operon. The *bgl* assay was performed as described above.

To determine the degree of functional similarity between GapR and H-NS, we constructed chimeras and compared them with the wild-type H-NS and GapR proteins with respect to their ability to repress gene expression in a Δ*h-ns E. coli* background. We measured mRNA levels of a subset of H-NS-repressed genes and tested the capability of cells to utilize the β-glucoside salicin, which depends on the expression of the *bgl* operon (29). Expression of H-NS, but not GapR, was found to reduce transcript levels of all H-NS-dependent genes analyzed and prevent salicin utilization (Fig. 1B, constructs 1 and 2). The coiled-coil motif in dimerization site 1 is necessary for H-NS to repress expression of the *bgl* operon and prevent salicin utilization (Fig. 1C, compare H-NS_Δ23-50_ and H-NS_Δ23-84_ with H-NS_1-137_). Fig. 1B shows that cells expressing chimeras in which the N-terminal region of GapR (residues 1-49) replaced either the H-NS dimerization site 1 and 2 (construct 3) or just dimerization site 1 (construct 4), failed to repress transcript levels of the H-NS-dependent genes and salicin utilization. However, as shown in Fig. 1B (constructs 5 and 6), expression of H-NS proteins containing a segment of GapR that comprises its coiled-coil motif (residues 20-47, Helix 1) fused to the N-terminal region of H-NS (residues 1-22) to create a chimeric dimerization site 1 led to decreased expression of all but one gene (*proV*) and compromised the ability of cells to grow in the presence of salicin. These results indicate that the N-terminal coiled-coil motif of GapR is functionally similar to the coiled-coil of H-NS, which is part of dimerization site 1. Interestingly, dimerization site 2 was found to be dispensable for both H-NS (Fig. 1C, H-NS_Δ51-84_) and a chimera containing the coiled-coil motif of GapR (Fig. 1B, construct 5) to silence gene expression, implying that the assembly of H-NS into higher order structures is not critical for repression of the H-NS-dependent genes tested with the exception of *proV*. We observed that the chimera H-NS_1-22_-GapR_20-49_-H-NS_85-137_ can only partially prevent salicin utilization (Fig. 1B, right panel, construct 5). However, no significant difference between construct 5 (H-NS_1-22_-GapR_20-49_-H-NS_85-137_) and construct 6 (H-NS_1-22_-GapR_20-47_-H-NS_51-137_) was observed upon measuring the transcript levels of H-NS-dependent genes in cells grown in the presence of the inducer for the prolonged period used in the assay (Fig. 1B left panel). Thus, the N-terminal coiled-coil motif of GapR can functionally replace the H-NS N-terminal coiled-coil motif.

Repression of the *bgl* operon has been shown to occur in *E coli* strains carrying only an H-NS truncated protein in which the entire DNA-binding domain is absent (30). We confirmed this finding by monitoring salicin utilization and the transcript levels of the *bgl* operon in a strain carrying an H-NS protein containing only the oligomerization domain (Fig. 1B, construct 7). Although *bglG* and *csgD* were both significantly repressed by H-NS lacking the entire DNA binding domain, the expressions of *gadB*, and o*mpC* were not repressed, and *proV* exhibited minimal repression (Fig. 1B left panel, construct 7). Robust repression of *gadB* and *proV* was rescued by a chimera in which Helix 2 and Helix 3 of GapR (residues 50-89) replaced part of the DNA-binding motif of H-NS (construct 9). We note that neither *gadB* nor *proV* were significantly repressed in construct 8, which has the entire H-NS C-terminal DNA-binding domain replaced by Helix 2 and Helix 3 of GapR (Fig. 1B left panel). Based on these results, we propose that, in addition to the shared function of their N-terminal coiled-coil motif, GapR and H-NS also share a functionally similar region in their C-termini.

### N-terminus of GapR drives self-association

Both GapR and H-NS share a coiled-coil motif at their N-termini (Fig. 1A). To determine if the N-terminus of GapR drives self-association, as does the corresponding region of H-NS (24–26), we used a bacterial two-hybrid assay and found that both a truncated WT protein comprising only Helix 1 (GapR_1-52_) and a full length WT protein (GapR_1-89_) interact with full-length GapR (Fig. 2A). Provided that GapR self-associates in a coiled-coil structure, oligomerization would be prevented by mutations replacing residues at positions “a” or “d” of the heptad repeats of the GapR coiled-coil motif. To test this hypothesis, the coding sequence corresponding to GapR_1-52_ was randomly mutagenized by error-prone PCR, and the amplicons were used to construct a library into the bacterial two-hybrid system. We identified three mutant *gapR*_1-52_ alleles (M1-M3) defective in oligomerization (Fig 2A). M1, M2 and M3 code for proteins replacing a residue at the position “a” or “d”. Curiously, when we carried out the bacterial two-hybrid assay with full length GapR protein containing the M1 and M3 alleles, we observed no defect in oligomerization (Fig. 2A). This result suggests that the presence of the DNA binding domain on the M1 and M3 GapR proteins stabilizes the association between the mutant and wild-type protein. The M2 allele, with the Q19R,L30P amino acid substitutions, in contrast, affected GapR self-association even when the C-terminus was present (Fig. 2A), indicating a more drastic consequence for the protein structure compared with the mutations found in the other alleles.

**Figure 2.**
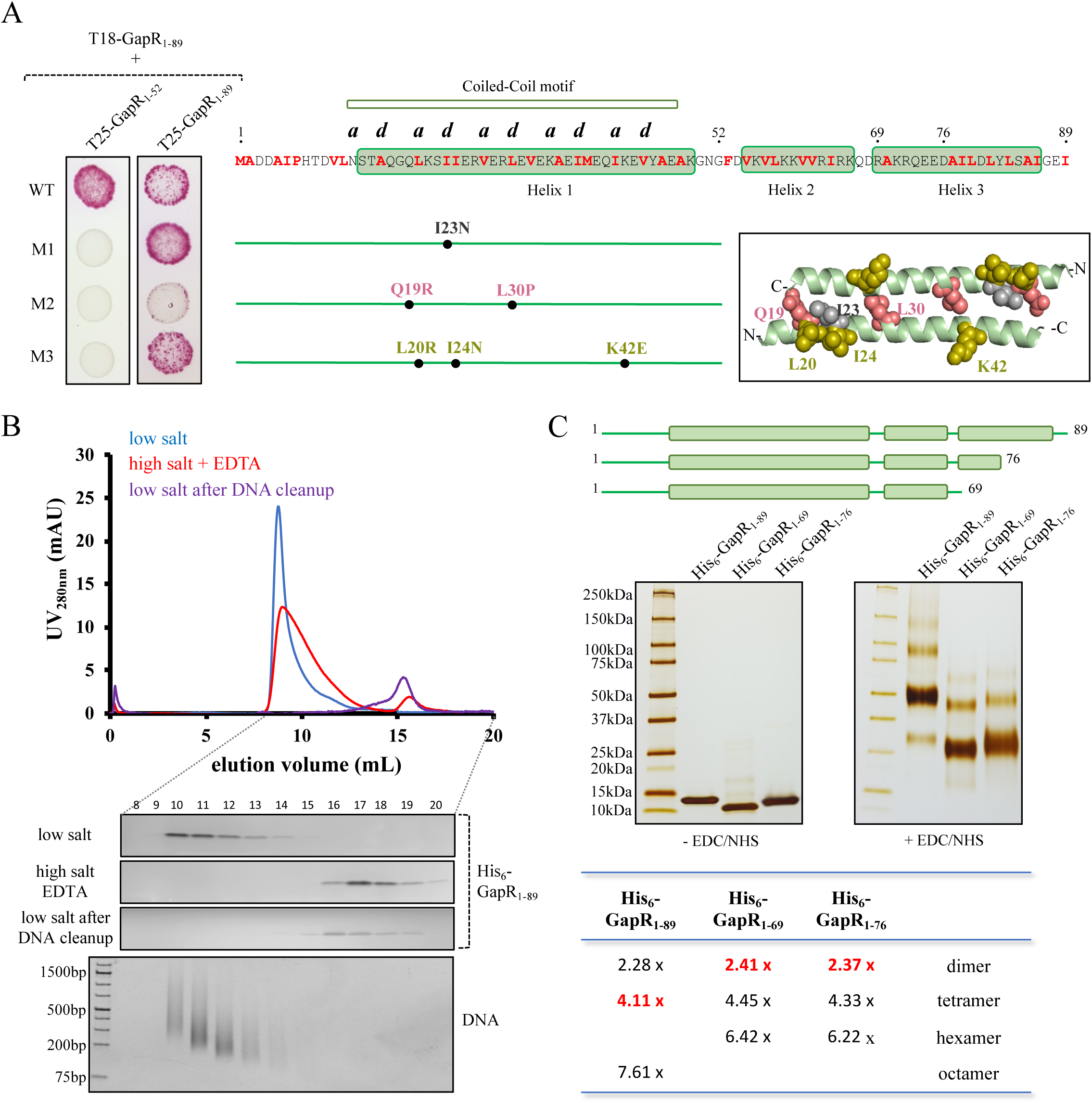
GapR forms a tetrameric structure. **A)** Screening for mutations that affect GapR self-association using a bacterial two-hybrid system. The left panel shows *E coli* BTH101 cells expressing the T18 domain of adenylate cyclase N-terminally fused to full-length WT GapR and the T25 domain of the same enzyme fused to the truncated N-terminus (GapR_1-52_ and the mutant GapR proteins M1, M2, or M3, isolated by random mutagenesis using error-prone PCR). Fusions of the T25 domain to full-length proteins (GapR_1-89_ and the mutant GapR proteins M1, M2, or M3) were also expressed in cells producing T18-GapR_1-89_. WT and mutant strains were grown to exponential phase (OD_600 nm_ of 0.5) and 3 µL from each culture were spotted in 1% maltose-containing MacConkey plates. The plates were imaged after 2 days at 30°C. The amino acid substitutions identified in the mutant alleles are shown in the middle panel. The predicted amino acid residues important for stabilization of canonical coiled-coil structures (denoted as “a” and “d”) and hydrophobic residues (highlighted in red) are indicated in the GapR sequence shown above the amino acid substitution mutants. The right panel shows the location of the native residues which were mutated in the M1, M2, and M3 alleles (shown as grey, salmon, and gold spheres, respectively) in the N-terminal GapR dimer derived from the DNA-bound crystal structure (PDB 6CG8) (15). **B)** Analysis of the full-length GapR by size exclusion chromatography. His_6_-GapR_1-89_ purified without nuclease treatment was dialyzed against buffer containing either 150 mM NaCl (low salt) or 1M NaCl supplemented with 1 mM EDTA (high salt + EDTA) and analyzed by size exclusion chromatography using the Superdex 200 10/300 GL column. His_6_-GapR_1-89_ separated from co-purified DNA was also analyzed by size exclusion chromatography (low salt after DNA cleanup). 50 µM protein (monomer) was used for all runs. Fractions 8-20 were resolved by SDS-PAGE, and the gels were silver stained. Shown below is a gel used to detect DNA prepared with fractions collected from protein treated with high salt + EDTA conditions and stained with ethidium bromide. **C)** Determination of the oligomerization state of WT GapR and two truncated GapR proteins. Proteins separated from co-purified DNA at 50 µM (monomer) were treated with crosslinking agents (400 mM EDC + 100 mM NHS) for 2 h at room temperature, the reaction products were resolved by SDS-PAGE, and the gel was silver stained. EDC and HNS act by a two-step reaction to crosslink glutamic and aspartic acid to lysine residue (50). Reactions were conducted in the presence of 1M NaCl and 1 mM EDTA as His_6_-GapR_1-69_ and His_6_-GapR_1-76_ precipitate in the presence of 150 mM NaCl. Under these conditions, GapR is not associated with DNA, ruling out the possibility that DNA could affect the oligomeric state of the proteins. As a control, proteins incubated for 2 h at room temperature in the absence of the crosslinking agents EDC/NHS were resolved by SDS-PAGE. The apparent molecular weight of each band was estimated and divided by the molecular weight calculated for the monomeric state of the corresponding protein. Values denoted in red refer to the main bands detected in the gel.

### GapR contains two non-contiguous dimerization sites that drive the assembly of a tetrameric structure

Affinity-purified full-length WT GapR_1-89_ was analyzed by size-exclusion chromatography. The protein eluted at the void volume of the Superdex 200 size exclusion column under low salt conditions (Fig. 2B), indicating soluble particles with a mass above 600 kDa. Upon treating the protein with a combination of high salt and EDTA, we observed a larger right-hand shoulder of the void peak corresponding to native DNA from the expression host and an additional elution peak far later than the void volume corresponding to GapR_1-89_ (Fig. 2B). These data suggest that GapR_1-89_ remains as nucleoprotein complexes throughout purification but dissociates into smaller soluble units when treated with high salt and EDTA. Because the smaller units of GapR_1-89_ that are separated from co-purified DNA in the presence of high salt and EDTA are no longer found in the void volume after decreasing salt and removing EDTA (Fig. 2B), we reasoned that DNA may act as a platform for the formation of higher order structures. Using a combination of size-exclusion chromatography and crosslinking experiments, both performed under high salt and EDTA conditions to prevent association of GapR with any contaminating DNA molecules, we determined that GapR_1-89_ alone is sufficient to assemble into a tetrameric structure (Fig. 2C and Fig. S1A), in agreement with structural data (15). We also analyzed untagged GapR_1-89_ and observed the same tetrameric state as the His-tagged protein (Fig. S1B). In crosslinking experiments, faint bands were observed in addition to the main band corresponding to tetrameric GapR (4.11-fold the molecular weight of the monomer) (Fig. 2C). The faint bands may correspond to reaction products in which only two out of the four subunits were crosslinked (2.28-fold the molecular weight of the monomer) or to nonspecific crosslinking of tetramers (7.61-fold the molecular weight of the monomer) (see the table at the bottom on Fig. 2C). Alternatively, these bands could indicate the presence of GapR at oligomeric states other than tetramer, which cannot be detected by size-exclusion chromatography (Fig. S1A).

Apart from the coiled-coil motif, GapR has two additional α-helices (Helix 2 and Helix 3). Helix 3 contains a cluster of hydrophobic residues at its C-terminal end (Fig. 2A). To determine the contribution of this region to oligomerization, we constructed truncated proteins lacking either the entire Helix 3 (GapR_1-69_) or only the hydrophobic patch in this helix (GapR_1-76_). These mutations caused a switch from the tetrameric state observed for GapR_1-89_ to a dimeric state as determined for both GapR_1-69_ and GapR_1-76_ (Fig. 2C and Fig. S1A). Again, according to the calculated molecular weight shown in the table at the bottom on Fig. 2C, the faint bands may represent nonspecific crosslinking of dimers or the ability of the truncated proteins GapR_1-69_ and GapR_1-69_ to form higher order structures. This result argues that Helix 3 in the C-terminal region of GapR represents a critical structural element for assembling the protein into a tetramer, in agreement with interactions illustrated by the DNA-bound crystal structure of GapR (15). Cumulatively, our data provides evidence that GapR forms a tetrameric structure by means of two non-contiguous dimerization sites (Helix 1 and Helix 3) and explains why a mutant of GapR_1-89_ carrying amino acid substitutions in its coiled-coil motif is still able to self-associate.

### GapR oligomerization is curtailed upon disruption of secondary structure

We described above the isolation of a *gapR* allele (M2, Fig. 2A) that encodes a protein carrying the Q19R,L30P amino acid substitutions in its coiled-coil motif and found that it exhibited defective oligomerization. Although the full-length GapR bearing these substitutions showed decreased protein solubility, the GapR_1-52_/Q19R,L30P truncation was expressed and purified as a soluble protein (Fig. S2). In crosslinking experiments, GapR_1-52_/Q19R,L30P was identified as a monomer (Fig. 3A), confirming that the Q19R,L30P mutations curtail GapR oligomerization. Differing elution volumes of the two proteins subjected to size exclusion chromatography corroborates this finding (Fig. S3). We carried out small angle X-ray scattering (SAXS) measurements of both GapR_1-52_ and GapR_1-52_/Q19R,L30P. The SAXS profiles were significantly different between the WT and mutant proteins and confirmed that the proteins do not form aggregates in solution (Fig. 3B). Guinier analysis yielded similar radii of gyration (Rg) for the wild-type and the mutant proteins (Fig. 3B), suggesting that despite the disruption in oligomeric state, the spherical dimensions of the soluble protein particles created by GapR_1-52_ and GapR_1-52_/Q19R,L30P are roughly the same. The pair-distance distribution functions of the SAXS profiles display a positive skew (Fig. 3C), suggesting that both proteins adopt elongated structures. Interestingly, the Kratky plots, which give us information about the flexibility and folding state of the proteins (31), differ significantly starting at mid-q values (∼0.05 1/Å) (Fig. 3D), with GapR_1-52_/Q19R,L30P failing to return to baseline at high q values. Therefore, GapR_1-52_/Q19R,L30P is more flexible than GapR_1-52_, raising the possibility of a difference in folded state. Circular dichroism (CD) analysis revealed that the fold of GapR_1-52_/Q19R,L30P is partially compromised when compared with GapR_1-52_ (Fig. 3E). Taken together, our results show that the helical secondary structure of the N-terminus of GapR is unfolded by the Q19R,L30P mutations and that this structural change precludes oligomerization.

**Figure 3.**
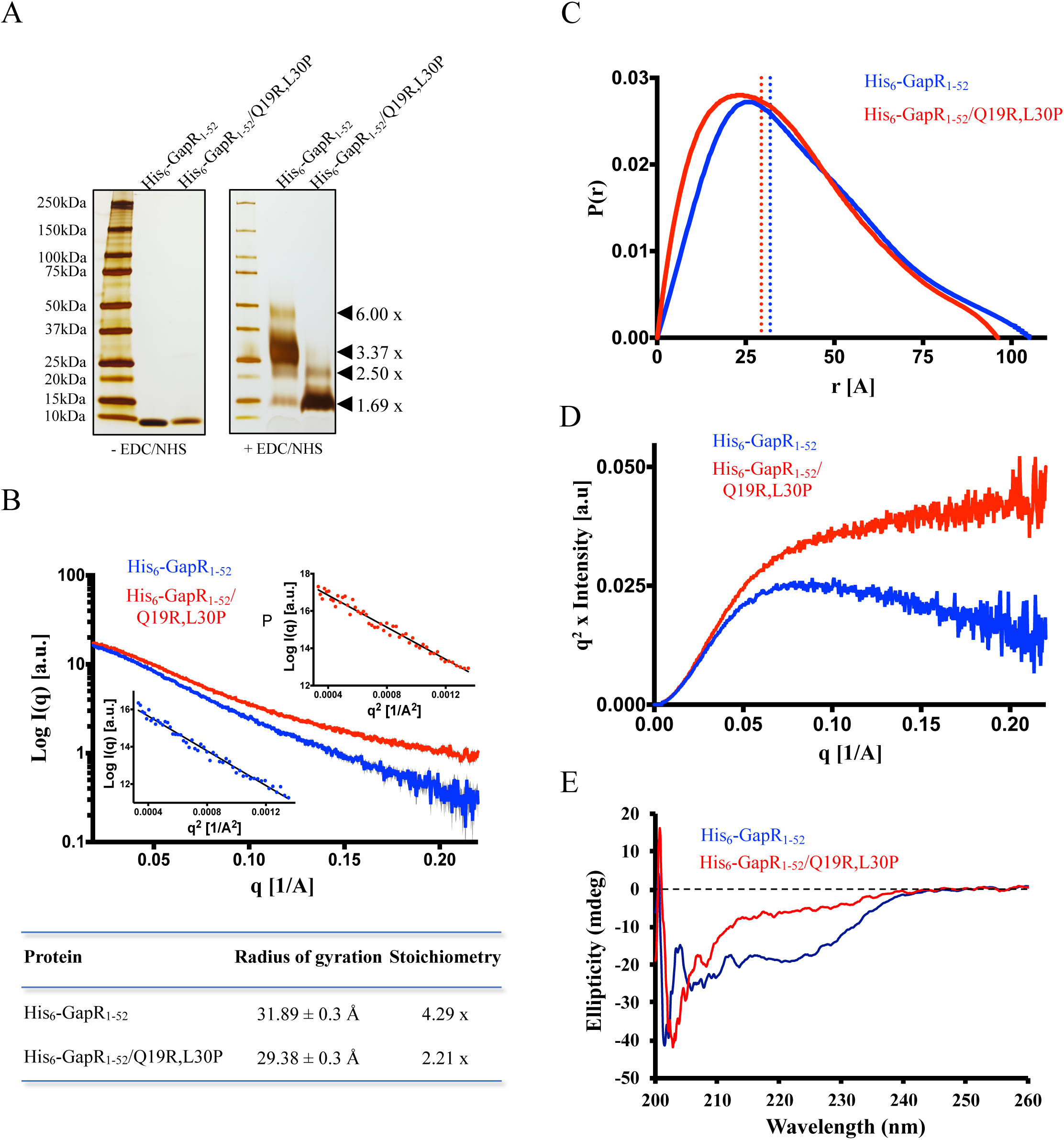
The N-terminal domain of GapR bearing the Q19R,L30P amino acid substitutions in the coiled-coil motif exhibits altered folding, flexibility and oligomeric state. **A)** Determination of the oligomerization state of truncated GapR proteins. Proteins at 50 µM (monomer) were crosslinked using 400 mM EDC + 100 mM NHS for 2 h at room temperature, the reaction products were resolved by SDS-PAGE, and the gel was silver stained. Reactions were conducted in the presence of 150 mM NaCl. As a control, proteins incubated for 2 h at room temperature in the absence of the crosslinking agents were resolved by SDS-PAGE. The apparent molecular weight of each band was estimated and divided by the molecular weight calculated for the monomeric state of the corresponding protein. **B-D)** SAXS analysis of truncated GapR proteins at 500 µM (monomer). **B)** SAXS profiles of the proteins. Insets: Guinier regimes used to calculate Rg of each protein. **C)** Pair-distance distribution functions for the proteins. Rg values are noted by vertical dotted lines. **D)** Kratky plots for the proteins. **E)** Circular Dichroism of truncated GapR proteins at 25 µM (monomer).

By using two-step homologous recombination, we replaced the *gapR* coding sequence with the mutant Q19R,L30P allele. We observed that cell growth was compromised at both 22° and 30°C and the cells exhibited a cell division defect, with a significant increase in the mean cell length and a broader distribution of sizes (Fig. S4 and Table S3).

### Positively charged amino acid residues at the C-terminus are critical for the DNA-binding activity of GapR

As shown above, GapR and H-NS share a second functionally similar region: Helix 2 of GapR (14) and a segment of the C-terminal DNA-binding domain of H-NS (27). To confirm that this region of GapR is critical for binding DNA, we first compared the truncated GapR_1-52_ and the full length GapR_1-89_ with respect to their ability to co-purify with and directly bind DNA *in vitro*. While the strong association of GapR_1-89_ with DNA accounts for the detection of the protein as part of nucleoprotein complexes in the void volume of the Superdex 200 size-exclusion chromatography column, GapR_1-52_ was retarded in its passage through the column, and no detectable amount of DNA was found to be associated with the protein (Fig. S5). Furthermore, electrophoretic mobility shift assay (EMSA) showed that the removal of most co-purified DNA from GapR_1-89_ allowed the protein to directly bind and shift the electrophoretic migration of the *pilA* promoter DNA fragment (Fig. 4A upper panel), a locus previously identified as a target of GapR *in vivo* (16). The absence of the C-terminal region in GapR_1-52_, in contrast, led to a protein completely deficient in binding DNA (Fig. 4A upper panel). Taken together, our data confirm that the C-terminus of GapR is necessary for the DNA-binding activity of this protein.

**Figure 4.**
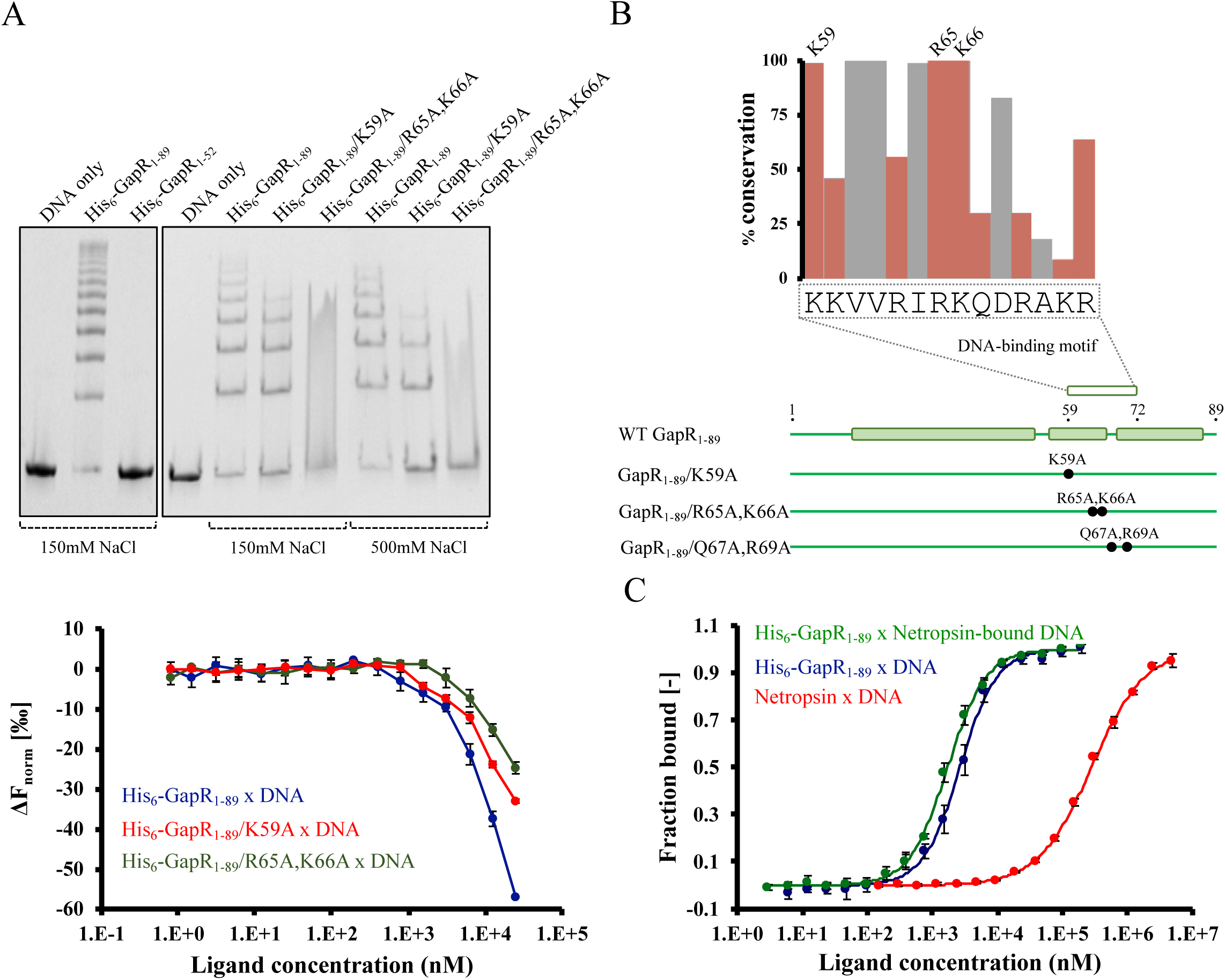
GapR binds DNA using highly conserved, positively charged residues at its C-terminal region and does not selectively recognize the narrow minor grove. **A)** Analysis of the DNA-binding affinity of WT and mutant GapR proteins by electrophoretic mobility shift assays (top) and microscale thermophoresis (bottom). For the electrophoretic mobility shift assay, 2.5 µM protein (monomer) was incubated with 0.1 µM 320 bp P_pilA_ DNA for 30 min at room temperature in the presence of either 150 or 500 mM NaCl, the reaction products were resolved by PAGE under native conditions, and the gels were stained with ethidium bromide. Controls with no protein included are also shown. For the microscale thermophoresis experiments, 0.1 µM ATTO 488-labelled 320 bp P_pilA_ DNA fragment was incubated with WT and mutant GapR proteins (0.8 nM to 25.0 µM) for 30 min at room temperature in the presence of 150 mM NaCl, and thermophoresis was determined for each sample. Values are changes in the normalized fluorescence upon heating (ΔF_norm_ = F_hot_/F_cold_) from 3 independent measurements. **B)** Conservation of amino acid residues in the predicted DNA-binding motif of GapR. The deduced amino acid sequence of 101 GapR orthologs was globally aligned using Clustal Omega (48), and the conservation of positively charged (red) and hydrophobic (gray) residues in the predicted DNA-binding motif of *C. crescentus* GapR was determined. Even though a glutamine is present at position 67 of *C. crescentus* GapR, a positively charged residue is located at the corresponding position of several GapR orthologs. Only the predicted DNA-binding motif of GapR (14) is represented. **C)** Evaluation of the effect of Netropsin on the association of GapR with DNA by microscale thermophoresis. 0.1 µM ATTO 488-labelled 12-mer DNA fragment was treated, or not, with 1.25 µM Netropsin for 30 min at room temperature in the presence of 150 mM NaCl, GapR at varied concentrations (3.0 nM to 200 µM) was added, and the resulting mixtures were incubated for an additional 30 min at room temperature. DNA thermophoresis was determined for each sample. The concentration of Netropsin used in the assays was determined by incubating 0.1 µM ATTO 488-labelled 12-mer DNA fragment with different amount of Netropsin (0.15 µM to 5 mM) for 30 min at room temperature and monitoring DNA thermophoresis. Because DNA saturation was reached in these experiments, the MST data were represented as the fraction of bound DNA molecules at each concentration of the ligand. Results are from 3 independent measurements.

We sought to identify specific amino acid residues in the DNA-binding motif of GapR that are important for the association of GapR with DNA. An alignment of the deduced protein sequence of 101 GapR orthologs within the α Proteobacteria, while avoiding overrepresentation of species clustered to the same genus, allowed us to identify three highly conserved, positively charged residues in Helix 2 of GapR (K59, R65 and K66 in the *C. crescentus* protein) (Fig. 4B). To evaluate the role of these residues on the DNA-binding affinity of GapR, we carried out site-directed mutagenesis of *gapR* and purified two full-length mutant proteins, GapR_1-89_/K59A and GapR_1-89_/R65A,K66A. Compared with wild-type GapR_1-89_, GapR_1-89_/K59A was clearly compromised with respect to its capability to shift the electrophoretic mobility of the *pilA* promoter DNA fragment, especially under high ionic strength (Fig. 4A top panel). Because the nucleoprotein complexes containing GapR_1-89_/R65A,K66A form a smear rather than individual bands in EMSA assay (Fig. 4A upper panel), a precise comparison between this mutant protein and WT GapR was not possible using this assay. To circumvent this problem, we compared the DNA-binding activity of the GapR proteins using microscale thermophoresis (MST) (32), which measured the change in the thermophoresis of the fluorescently-labeled *pilA* promoter DNA fragment caused by the binding of GapR proteins. As we could not obtain GapR_1-89_/R65A,K66A at concentrations higher than 25 μM, MST experiments were carried out with all proteins up to 25 μM. Even though these protein concentrations were not sufficient to reach DNA saturation and determine the dissociation constants, we observed that the mutant proteins at any concentration ranging from 0.8 to 25 μM led to smaller changes in the normalized fluorescence of DNA molecules upon heating (ΔF_norm_) when compared with WT GapR_1-89_ (Fig. 4A bottom panel). This data implies decreased binding of the mutant proteins to DNA relative to WT GapR. Further, we showed that GapR_1-89_/R65A,K66A affected the variation of the normalized fluorescence of DNA molecules to a lesser extent compared with GapR_1-89_/K59A (Fig. 4A bottom panel), indicating a more severe effect of substituting R65 and K66 relative to the K59A mutation. Strikingly, none of the point mutations analyzed completely disrupted DNA-binding (Fig. 4A bottom panel), indicating that more than one protein-DNA contact is critical for stabilizing the nucleoprotein complexes.

By replacing the *gapR* coding sequence with the K59 and R65A,K66A alleles (Fig. S4A), we observed that the growth of cells expressing the mutant allele corresponding to GapR_1-89_/R65A,K66A was affected at 30°C, while cells containing the K59A mutation exhibited no growth defect (Fig. S4B). Although the K59A mutation slightly increased the mean cell length, strains carrying the R65A,K66A allele in place of wild type *gapR* exhibited aberrant cell division with a significant increase in the mean cell length (Fig. S4C and Table S3). Curiously, cells carrying the R65A,K66A mutation were more elongated at 30°C than at 22°C (Fig. S4C and Table S3), corroborating our finding that the growth of these cells is compromised only at 30°C. Thus, mutations disturbing GapR function by reducing DNA-binding compromise cell fitness. Consistent with the hypothesis that amino acid residues contacting DNA are conserved throughout GapR orthologs, the substitution of two non-conserved, polar residues (Q67 and R69) caused no effect on the DNA-binding activity of GapR when the binding event was monitored in the presence of low salt and only a minor effect under high ionic strength (Fig. S6). Based on the subdomain organization of GapR, we predicted that the K59A, R65A,K66A and Q67A,R69A mutations would have no effect on oligomerization. Indeed, interaction of the mutant proteins with the wild-type protein was quite similar to the self-association of wild-type GapR (Fig. S7). Therefore, our analyses identified highly conserved, positively charged amino acid residues in the DNA-binding motif of GapR that are specifically important for the DNA-binding activity of the protein.

### GapR does not selectively recognize a narrow minor groove at AT-rich DNA loci

Several NAPs achieve selective binding to AT-rich DNA loci by inserting specific polar amino acids into the narrow minor groove characteristic of these sequences (33). Highly conserved, positively charged amino acid residues are important for the DNA-binding activity of GapR, implying that a different mechanism may contribute to this binding specificity. To explore this possibility, we determined the effect of the minor groove binding reagent Netropsin on the affinity of GapR_1-89_ for a 12-mer DNA fragment containing the (dAT)_3_ sequence. Netropsin was previously shown to bind to this DNA sequence and to compete with H-NS binding (34). Upon monitoring thermophoresis, we confirmed the association of Netropsin with the 12-mer DNA substrate and found that Netropsin does not compete for the binding of GapR_1-89_ to DNA (Fig. 4C). Instead, the binding affinity of GapR_1-89_ for Netropsin-bound 12-mer DNA (EC_50_ = 1.73 ± 0.05 µM) is slightly higher than that determined for the interaction of GapR_1-89_ with DNA (EC_50_ = 2.67 ± 0.17 µM) (Fig. 4C). We also analyzed the effect of Netropsin on the electrophoretic migration of the *pilA* promoter DNA and found that Netropsin changes the migration profile of the DNA, confirming binding of Netropsin to a DNA fragment with an overall AT content of 53% (Fig. S8). In addition, at the highest Netropsin concentration used, GapR_1-89_ had a more drastic effect on DNA retardation compared with a sample with no Netropsin added (Fig. S8). Therefore, the results from these competition experiments indicate that GapR specifically recognizes AT-rich DNA sequences by a mechanism other than direct binding to narrow minor groves, which agrees with the finding that the maximum DNA-binding activity of GapR depends on highly conserved, positively charged amino acid residues.

### GapR stimulates DNA bridging *in vitro*

To determine the effect of GapR binding on the nanoscale organization of DNA, we visualized single nucleoprotein complexes with fluid-atomic force microscopy (AFM). Fluid-AFM provides sufficient spatial resolution to observe molecular sub-structures without fixing or drying biological samples, thus preserving their native state (35). AFM of DNA molecules containing the *Caulobacter pilA* promoter showed well-separated single molecules that matched the expected length (∼340 nm) (Fig. 5A top, left panel). Random overlaps within DNA molecules created a low percentage of intramolecular junctions. However, the addition of GapR_1-89_ to DNA, when imaged under the same conditions as DNA alone, displayed a higher frequency of intramolecular junctions compared to DNA molecules incubated with no protein (Fig. 5A top, middle panel). This increase in intramolecular junctions was lost when DNA was incubated with the purified mutant GapR protein that is defective in DNA binding (Fig. 5A top, right panel). Upon increasing the sample density, we observed an increase in intermolecular junctions with the nucleoprotein complex compared to DNA alone (Fig. 5B). AFM is capable of providing the relative heights of features in a given sample, which serves as a proxy for presence or absence of proteins for both intramolecular and intermolecular junctions. Accordingly, we compared feature heights at both junction and off-junction regions in the presence or absence of the GapR protein (Fig. 5A bottom panel and Fig. 5B bottom panel). Quantification of junction height differences showed that the relative increase in height at the junctions in DNA molecules incubated with GapR_1-89_ was significantly greater than those measured for DNA molecules without GapR_1-89_ (Fig. 5A bottom panel, Fig. 5B bottom panel and Fig. 5C). As expected, a mutant GapR protein defective in DNA binding, GapR_1-89_/R65A,K66A, exhibited similar height differences as observed at random overlaps with DNA alone (Fig. 5C). Therefore, the increase in junction heights can be attributed to the presence of GapR and its association with DNA at these junctions. We asked whether GapR binds non-specifically to DNA junctions, or if GapR affects the formation of these junctions? To answer this question, we quantified junctions across DNA, DNA-GapR_1-89_ and DNA-GapR_1-89_/R65A,K66A samples as a percentage of total observed molecules. To rule out density dependent effects, we performed these measurements and analyses at two different densities of molecules in a 1 µm^2^ field of view (Fig. 5D). We found a significantly higher percentage of junctions in DNA molecules when GapR_1-89_ was present compared with DNA molecules imaged in the absence of the protein, both at high and low molecular density (Fig. 5D). In agreement with a decreased binding affinity of GapR_1-89_/R65A,K66A for DNA (Fig. 4A) and a smaller junction height difference observed for this protein (Fig. 5C), the DNA-GapR_1-89_/R65A,K66A sample showed a percentage of junctions similar to that calculated for DNA alone (Fig. 5D). We further asked whether GapR has a preference for inter- or intramolecular junctions. Upon doubling the molecular density in a field of view, we observed a 25-fold increase in the percentage of intermolecular junctions, while the percentage of intramolecular junctions was reduced by 1.5-fold (Fig. 5E). This result suggests that GapR does not have a preference for inter- or intramolecular association *in vitro*. We also noted that the junctions formed by GapR at the tested concentrations do not spread as those observed with H-NS nucleoprotein complexes using AFM (8). Taken together, the AFM experiments show that GapR can stimulate inter- and intramolecular bridging of DNA *in vitro*.

**Figure 5.**
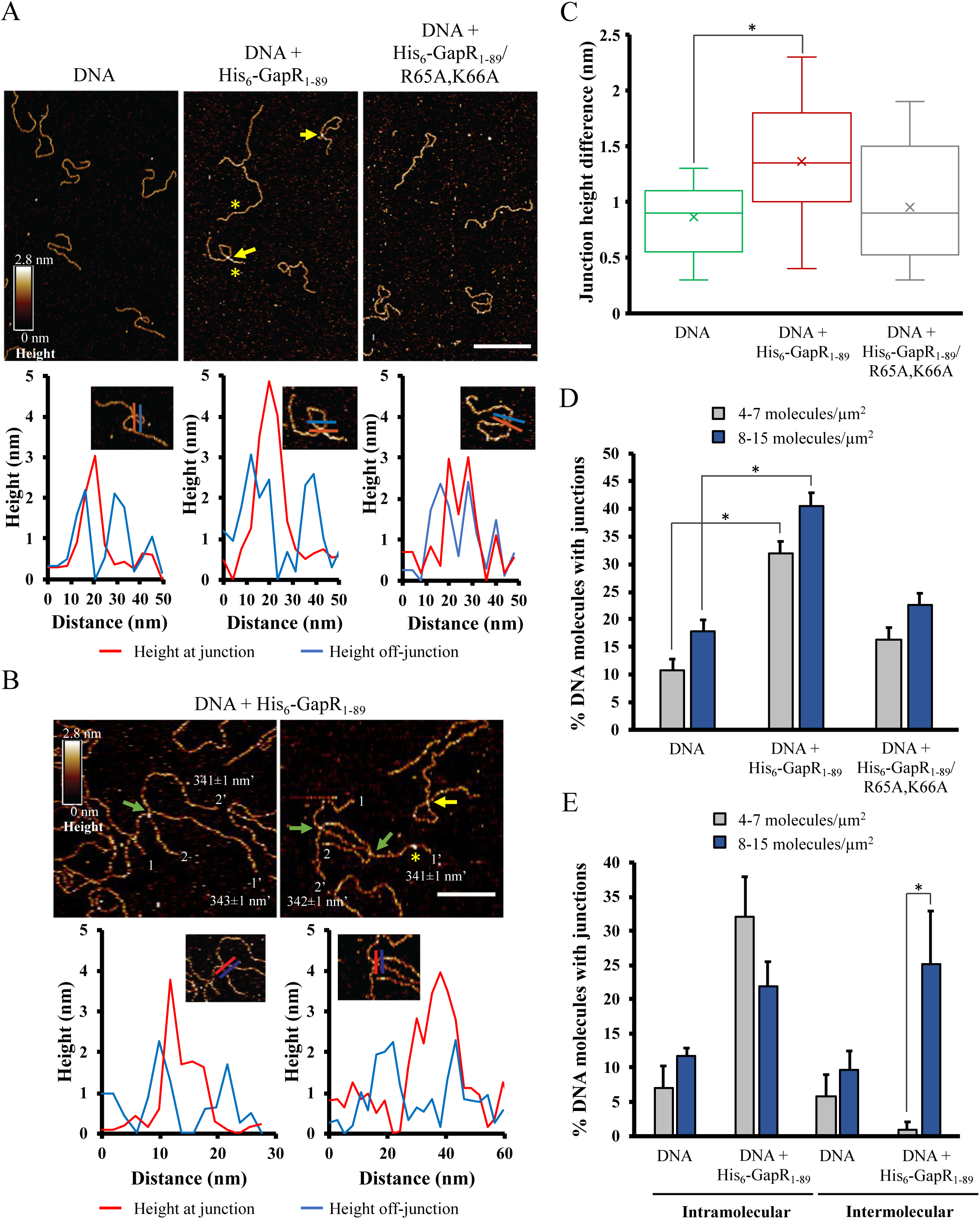
GapR binding stimulates the formation of junctions in DNA molecules. **A)** Fluid atomic force microscopy scans of a 1 kb DNA fragment with no protein added, incubated with 100 nM GapR_1-89_ (WT) or 100 nM GapR_1-89_/R65A,K66A (mutant defective in DNA-binding). Line profiles showing the height relative to the mica substrate for selected DNA molecules are shown below each image. The height at the junction (red line) was compared with the height away from the junction (blue lines). Yellow arrows indicate junctions with increased height of GapR-DNA junctions compared with junctions observed in the DNA molecules incubated with no protein added. Yellow asterisks indicate likely observations of GapR-DNA complexes (assessed from heights). **B)** Representative images of 1kb DNA incubated with 100 nM GapR_1-89_ and imaged using fluid AFM show molecules with intermolecular junction (green arrows) or intramolecular junction (yellow arrow). Yellow asterisk indicates likely observation of a GapR-DNA complex (assessed from heights). The termini of each molecule are marked by the number 1 and 1’ (for molecule 1) and 2 and 2’ (for molecule 2). Next to the prime terminus, the average measured length of the molecule is shown from five repeats of manual measurement of the molecule using a freehand line selection tool in Fiji (51). Line profiles drawn over the junction and off-junction regions are shown in the panel below for each image for the region shown in the inset. **C)** Distribution of the junction height difference calculated for 17 junctions for DNA, 34 junctions for DNA + GapR_1-89_ and 12 junctions for DNA + GapR_1-89_/R65A,K66A. The junction height differences (nm), mean±SEM, were as follows: 0.86±0.08 nm (DNA with no protein added), 1.36±0.08 nm (DNA + GapR_1-89_) and 0.95±0.16 nm (DNA + GapR_1-89_/R65A,K66A). **D)** Quantification of junctions observed in each sample. These measurements were performed at two different densities of molecules. **E)** Bar plot showing the percentage of molecules of DNA (or DNA + GapR_1-89_) displaying intermolecular and intramolecular associations. These data are shown for both low (4-7 molecules/ µm^2^) and high (8-15 molecules/µm^2^) densities. Asterisk in bar plots represents statistically different values according to unpaired Student’s t-Test (p<0.001).

## DISCUSSION

### Structural bases for the similar and distinct functional aspects of GapR and H-NS

GapR and H-NS exhibit only short regions of sequence conservation and are quite different with respect to tertiary structure, oligomerization, and DNA-binding. However, GapR and H-NS share two functionally similar regions. One of these regions is Helix 1 of GapR, which forms a coiled-coil motif that drives self-association (Fig. 6A). GapR mutants with self-association defects disrupt the heptad repeats integral to forming a coiled-coil structure. The most severe of these mutants, Q19R, L30P, has Helix 1 unfolded and exhibits numerous growth defects, suggesting that robust oligomerization is directly linked to cellular fitness.

**Figure 6.**
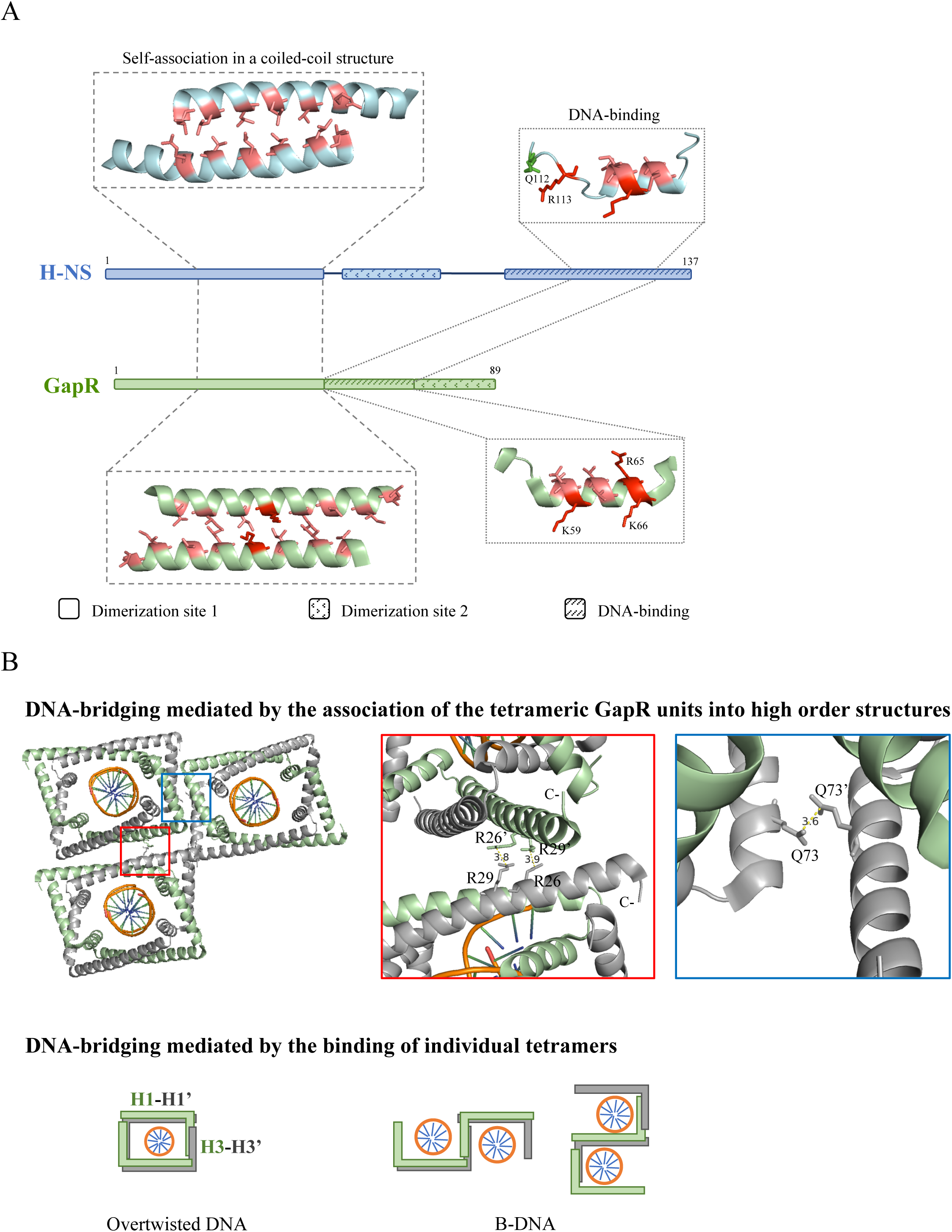
GapR shares with H-NS two structurally and functionally similar regions but bridges DNA differently. **A)** A schematic of the functional regions of GapR and H-NS highlighting the two conserved structural elements. The structural elements shown are part of the crystal structures of H-NS_1-83_ from *Salmonella enterica* serovar Typhimurium (PDB 3NR7) (24) and GapR_11-89_ from *Caulobacter crescentus* (PDB 6CG8) (15) and the solution structure of H-NS_91-137_ from *Escherichia coli* (PDB 1HNR) (27). Hydrophobic and positively charged residues are colored as salmon and red, respectively. Residues involved in DNA-binding (Q112 and R114 from H-NS and K59 and R66 from GapR) or playing a possible structural role (R65 from GapR) are indicated. **B**) Two potential modes of GapR-mediated DNA-bridging. Three DNA-bound GapR tetramers as observed in the crystal lattice of PDB 6CG8 (15) illustrate a possible model in which DNA-bridging is caused by the association of the tetrameric GapR units into high order structures. Two regions that may potentially mediate DNA-bridging interactions are included as zoom insets. The lower panel shows a schematic of the possible alternative conformations for B-DNA-bound GapR tetramer to illustrate the DNA-bridging model mediated by the binding of individual tetramers.

H-NS was previously structurally characterized by high-resolution methods including an X-ray crystal structure of the oligomerization domain, H-NS_1-83_, from *Salmonella enterica* serovar Typhimurium (24) and an NMR solution structure of the DNA-binding domain, H-NS_91-137_, from *Escherichia coli* (27). Aligning the H-NS DNA-binding domain to the crystal structure of the DNA-bound form of an H-NS related protein, Ler from enterohaemorrhagic *E. coli* (36), supports a model for full-length H-NS in which an H-NS oligomer simultaneously binds two segments of the nucleoid to form a superhelical nucleoprotein plectoneme (24). While self-association of H-NS via its coiled-coil region (dimerization site 1) is critical for gene silencing (24–26, 30, 37), other proteins have been identified that bind to and help H-NS modulate transcription, further complicating efforts to understand this small, multifunctional protein (38–40).

A chimera generated by replacing the H-NS coiled-coil motif with Helix 1 of GapR rescued gene regulation activity, highlighting the functional similarity between the two regions. However, the short N-terminal region (residues 1-22) preceding the H-NS coiled-coil motif, which consists of two short helices, is not interchangeable (Fig. 1A). These two small helices in H-NS rest against the coiled-coil motif of its neighbor, potentially stabilizing this interface (24). In addition, the N-terminal region of H-NS encompassing residues 1-22 is crucial for gene silencing as a H-NS mutant carrying the R12H substitution was reported to be defective in both repressing gene expression (30) and interacting with Hha (38), a protein required for fully silencing transcription of a subset of the H-NS regulon (41, 42). Even though the C-terminal domain of H-NS is the primary region for DNA-binding (30, 33), the N-terminal basic residues R15 and R19 may weakly interact with DNA strands that run alongside the oligomerization domain in the superhelical plectoneme structural model (24). This putative interaction between a dimerization site and a companion DNA molecule may also contribute to the crucial role of this H-NS region in silencing gene expression (25).

The N-terminus of GapR preceding its coiled-coil motif (coincident with Helix 1) does not possess basic residues and is not conserved among GapR orthologs (15). Although the basic residues K34, K42, and K49 in GapR Helix 1 do not support the binding of Helix 1 to DNA (GapR_1-52_ does not bind DNA (Fig. 4A)), these residues in Helix 1 were found to contact phosphates of the bound DNA molecule (15) and may stabilize the Helix 2-mediated association of the full-length protein with DNA. Therefore, distantly related NAP dimerization domains may serve two conserved functions, self-association and support of DNA-binding.

The second region shared by GapR and H-NS corresponds to a DNA-binding motif in Helix 2 of GapR (Fig 1A) and a region within the DNA-binding motif of H-NS (Fig. 6A). GapR Helix 2, rests against its bound DNA strand with both protein and DNA helices in parallel orientation (15). K59 and K66 contact the DNA phosphate backbone while R65 seems to play a structural role based on its possible participation in an electrostatic interaction with E28 from the other subunit in the same dimer (15). Although we did not analyze the R65A and K66A mutations individually, an E28A substitution slowed the migration of GapR under native PAGE without affecting oligomerization (Fig. S9). This effect is not caused by a change in the net charge of GapR, as an E31A mutation does not affect the electrophoretic migration of GapR (Fig. S9). Therefore, the E28A mutation likely leads to a structural change in the protein and the reduced DNA-binding activity of GapR/R65A,K66A is most probably the result of a combination of the loss of a residue directly contacting DNA and a residue playing a structural role. The involvement of positively charged residues in DNA-binding and our finding that GapR does not competitively bind the narrow minor groove of DNA agrees well with the reported crystal structure (15), which indicates that GapR binds to DNA mainly by interactions with the phosphate backbone. We observed that a chimera generated by fusing Helix 2 and Helix 3 of the DNA binding domain of GapR with the first two β-strands of the H-NS DNA-binding domain was able to silence gene expression. Therefore, it seems that a DNA-binding domain functionally similar to that found in the wild-type H-NS protein may result from the described chimeric construction. In this scenario, the helices would likely need to re-orient with respect to the DNA strand in order to maintain backbone phosphate interactions, shelter hydrophobic clusters, and keep polar residues exposed to solvent.

### Potential modes of DNA bridging by GapR

The conserved Helix 1 and Helix 2 in GapR leave its Helix 3 with no clear homology to H-NS (Fig 1A). We showed here that Helix 3 of GapR acts as a second dimerization site that is critical for the formation of the tetrameric state, as predicted by the analysis of the GapR crystal structure encircling DNA (15) (Fig, 6B). Using Fluid-AFM we observed that GapR can also mediate DNA bridging, possibly through self-association of the GapR-DNA complex (Fig 5). An inspection of the crystal structure of DNA-bound GapR (15) reveals that two arginine residues (R26 and R29) are located on the outer surface of GapR Helix 1 (Fig. 6B). Some of these clusters are in close proximity (3.5-4.0 Å apart) to pairs from a neighboring tetramer. Therefore, these residues have the potential to form an arginine cluster (43–46), which could mediate the association of the tetrameric GapR units into high order structures. Similarly, residue Q73 in GapR Helix 3 may form a close association (3.5-4.0 Å apart) with Q73 from a neighboring tetramer, offering an additional molecular interaction that may be involved in inter-tetramer associations (Fig. 6B).

Alternatively, we can envision a model in which DNA bridging is promoted by individual tetramers. In the recently reported crystal structure of GapR bound to an AT-rich DNA fragment (15), the DNA in an over-twisted state can be accommodated in the internal cavity of tetrameric GapR. While this structure provides the molecular basis for GapR binding to over-twisted DNA, it does not necessarily rule out the possibility that GapR may also bind relaxed B-DNA. In fact, the electrophoretic migration of a (dCG)_6_ DNA fragment is shifted by GapR *in vitro* (Fig. S10), demonstrating that GapR can indeed bind relaxed DNA. GapR likely cannot form a tetramer to clamp B-DNA due to the increased width of the DNA strand and GapR is not expected to over-twist DNA on its own (15). Therefore, the association of B-DNA with GapR may lead to an opened conformation of a tetrameric nucleoprotein assembly. Rotating one monomer with respect to its partner might also allow linked GapR subunits to bind more than one segment of the nucleoid simultaneously (Fig. 6B, lower panel).

DNA-bridging by GapR occurs *in vitro* as observed by fluid-AFM; however, while H-NS forms filaments in complex with DNA, GapR-DNA associations seem to occur at discrete, localized junctions. Both hypotheses presented above imply that GapR cannot form extended regions of bridged DNA, as the association of the tetramers to each other relies on relatively weak interactions (arginine clusters or single-residue interactions) or GapR binding to two B-DNA fragments may not be as stable as the interaction between GapR and over-twisted DNA. This limited bridging activity of GapR may also explain why transcription read-through is not favored in cells depleted of GapR (15), thus differentiating it from H-NS, which stimulates pausing and Rho-dependent termination (8). The biological significance of the DNA-bridging activity of GapR remains to be investigated in the context of the *Caulobacter* nucleoid organization.

### Evolutionary relationship of GapR and H-NS

The members of the α-Proteobacteria predominantly contain *gapR* and rarely contain *hns* (Fig. S11). Both of these nucleoid-associated proteins interact preferentially with DNA regions of high AT content and bridge DNA, while using different means of oligomerization to affect genome transcription. GapR forms a tetramer that encircles over-twisted DNA to enable transcription progression (15) while H-NS forms higher order oligomers that repress gene expression (47). The discovery that GapR and H-NS share two structurally and functionally similar regions (Fig. 1) implies an evolutionary relationship between these proteins. The simplest explanation is that *gapR* may have arisen very early after the α-Proteobacteria emerged as a separate phylogenetic group. Based on the primary structure comparison shown here (Fig. 1A), regions of H-NS are conserved in GapR, while the central part of H-NS is missing in GapR (Fig. 6A). Analysis of chimeras between the two proteins revealed that the evolutionarily conserved self-association coiled-coil region and a DNA binding region maintained function (Fig. 1 and Fig. 6A). Regions of H-NS that are either deleted or structurally distinct in GapR function to repress gene expression. The acquisition of sequences to the evolved GapR protein gave rise to the ability to form tetramers that encircle DNA (15), while retaining the inability to silence gene expression. Judging by the broad occurrence of *hns* in the α-proteobacterial family *Rhodobacterales* (Fig. S11), it is more parsimonious to assume that a bacterium ancestral to this family acquired *hns*, and this gene has been maintained in the majority of the actual species. In contrast, the sporadic presence of *hns* in another α-proteobacterial family, *Rhizobiales,* suggests multiple, more recent horizontal transfer events. Understanding why GapR has evolved and been maintained in the majority of the α-Proteobacteria, and H-NS is rarely present in this group, are critical points for future investigation.

## ACKNOWLEDGEMENTS

We thank Marcin Walkiewicz (Stanford Nano Shared Facility), Cedric Espenel (Cell Sciences Imaging Facility, Stanford University) and Pascal Odermatt (School of Dentistry, UCSF) for helpful discussions on fluid-AFM experiments. Use of the Stanford Synchrotron Radiation Lightsource, SLAC National Accelerator Laboratory, is supported by the U.S. Department of Energy, Office of Science, Office of Basic Energy Sciences under Contract No. DE-AC02-76SF00515. The SSRL Structural Molecular Biology Program is supported by the DOE Office of Biological and Environmental Research, and by the National Institutes of Health, National Institute of General Medical Sciences (including P41GM103393). The contents of this publication are solely the responsibility of the authors and do not necessarily represent the official views of NIGMS or NIH.

## REFERENCES

1. Wang, X., Montero Llopis, P. and Rudner, D.Z. (2013) Organization and segregation of bacterial chromosomes. Nat Rev Genet, 14, 191–203.

2. Dame, R.T. (2005) The role of nucleoid-associated proteins in the organization and compaction of bacterial chromatin. Mol Microbiol, 56, 858–870.

3. Grainger, D.C., Hurd, D., Goldberg, M.D. and Busby, S.J. (2006) Association of nucleoid proteins with coding and non-coding segments of the *Escherichia coli* genome. Nucleic Acids Res, 34, 4642–4652.

4. Kahramanoglou, C., Seshasayee, A.S., Prieto, A.I., Ibberson, D., Schmidt, S., Zimmermann, J., Benes, V., Fraser, G.M. and Luscombe, N.M. (2011) Direct and indirect effects of H-NS and Fis on global gene expression control in *Escherichia coli*. Nucleic Acids Res, 39, 2073–2091.

5. Lucchini, S., Rowley, G., Goldberg, M.D., Hurd, D., Harrison, M. and Hinton, J.C. (2006) H-NS mediates the silencing of laterally acquired genes in bacteria. PLoS Pathog, 2, e81.

6. Navarre, W.W., Porwollik, S., Wang, Y., McClelland, M., Rosen, H., Libby, S.J. and Fang, F.C. (2006) Selective silencing of foreign DNA with low GC content by the H-NS protein in *Salmonella*. Science, 313, 236–238.

7. Oshima, T., Ishikawa, S., Kurokawa, K., Aiba, H. and Ogasawara, N. (2006) *Escherichia coli* histone-like protein H-NS preferentially binds to horizontally acquired DNA in association with RNA polymerase. DNA Res, 13, 141–153.

8. Kotlajich, M.V., Hron, D.R., Boudreau, B.A., Sun, Z., Lyubchenko, Y.L. and Landick, R. (2015) Bridged filaments of histone-like nucleoid structuring protein pause RNA polymerase and aid termination in bacteria. Elife, 4.

9. Tendeng, C. and Bertin, P.N. (2003) H-NS in Gram-negative bacteria: a family of multifaceted proteins. Trends Microbiol, 11, 511–518.

10. Gordon, B.R., Imperial, R., Wang, L., Navarre, W.W. and Liu, J. (2008) Lsr2 of *Mycobacterium* represents a novel class of H-NS-like proteins. J Bacteriol, 190, 7052–7059.

11. Gordon, B.R., Li, Y., Wang, L., Sintsova, A., van Bakel, H., Tian, S., Navarre, W.W., Xia, B. and Liu, J. (2010) Lsr2 is a nucleoid-associated protein that targets AT-rich sequences and virulence genes in *Mycobacterium tuberculosis*. Proc Natl Acad Sci U S A, 107, 5154–5159.

12. Smits, W.K. and Grossman, A.D. (2010) The transcriptional regulator Rok binds A+T-rich DNA and is involved in repression of a mobile genetic element in *Bacillus subtilis*. PLoS Genet, 6, e1001207.

13. Chen, J.M., Ren, H., Shaw, J.E., Wang, Y.J., Li, M., Leung, A.S., Tran, V., Berbenetz, N.M., Kocincova, D., Yip, C.M. et al. (2008) Lsr2 of *Mycobacterium tuberculosis* is a DNA-bridging protein. Nucleic Acids Res, 36, 2123–2135.

14. Arias-Cartin, R., Dobihal, G.S., Campos, M., Surovtsev, I.V., Parry, B. and Jacobs-Wagner, C. (2017) Replication fork passage drives asymmetric dynamics of a critical nucleoid-associated protein in *Caulobacter*. EMBO J, 36, 301–318.

15. Guo, M.S., Haakonsen, D.L., Zeng, W., Schumacher, M.A. and Laub, M.T. (2018) A Bacterial Chromosome Structuring Protein Binds Overtwisted DNA to Stimulate Type II Topoisomerases and Enable DNA Replication. Cell, 175, 583–597 e523.

16. Ricci, D.P., Melfi, M.D., Lasker, K., Dill, D.L., McAdams, H.H. and Shapiro, L. (2016) Cell cycle progression in *Caulobacter* requires a nucleoid-associated protein with high AT sequence recognition. Proc Natl Acad Sci U S A, 113, E5952–E5961.

17. Taylor, J.A., Panis, G., Viollier, P.H. and Marczynski, G.T. (2017) A novel nucleoid-associated protein coordinates chromosome replication and chromosome partition. Nucleic Acids Res, 45, 8916–8929.

18. Gibson, D.G., Young, L., Chuang, R.Y., Venter, J.C., Hutchison, C.A., 3rd and Smith, H.O. (2009) Enzymatic assembly of DNA molecules up to several hundred kilobases. Nat Methods, 6, 343-345.

19. Cadwell, R.C. and Joyce, G.F. (1992) Randomization of genes by PCR mutagenesis. PCR Methods Appl, 2, 28–33.

20. Untergasser, A., Cutcutache, I., Koressaar, T., Ye, J., Faircloth, B.C., Remm, M. and Rozen, S.G. (2012) Primer3-new capabilities and interfaces. Nucleic Acids Res, 40, e115.

21. Schmittgen, T.D. and Livak, K.J. (2008) Analyzing real-time PCR data by the comparative C(T) method. Nat Protoc, 3, 1101–1108.

22. Schmittgen, T.D. (2001) Real-time quantitative PCR. Methods, 25, 383–385.

23. Kim, E.A. and Blair, D.F. (2015) Function of the Histone-Like Protein H-NS in Motility of *Escherichia coli*: Multiple Regulatory Roles Rather than Direct Action at the Flagellar Motor. J Bacteriol, 197, 3110–3120.

24. Arold, S.T., Leonard, P.G., Parkinson, G.N. and Ladbury, J.E. (2010) H-NS forms a superhelical protein scaffold for DNA condensation. Proc Natl Acad Sci U S A, 107, 15728–15732.

25. Bloch, V., Yang, Y., Margeat, E., Chavanieu, A., Auge, M.T., Robert, B., Arold, S., Rimsky, S. and Kochoyan, M. (2003) The H-NS dimerization domain defines a new fold contributing to DNA recognition. Nat Struct Biol, 10, 212–218.

26. Esposito, D., Petrovic, A., Harris, R., Ono, S., Eccleston, J.F., Mbabaali, A., Haq, I., Higgins, C.F., Hinton, J.C., Driscoll, P.C. et al. (2002) H-NS oligomerization domain structure reveals the mechanism for high order self-association of the intact protein. J Mol Biol, 324, 841–850.

27. Shindo, H., Iwaki, T., Ieda, R., Kurumizaka, H., Ueguchi, C., Mizuno, T., Morikawa, S., Nakamura, H. and Kuboniwa, H. (1995) Solution structure of the DNA binding domain of a nucleoid-associated protein, H-NS, from *Escherichia coli*. FEBS Lett, 360, 125–131.

28. Gao, Y., Foo, Y.H., Winardhi, R.S., Tang, Q., Yan, J. and Kenney, L.J. (2017) Charged residues in the H-NS linker drive DNA binding and gene silencing in single cells. Proc Natl Acad Sci U S A, 114, 12560–12565.

29. Prasad, I. and Schaefler, S. (1974) Regulation of the beta-glucoside system in *Escherchia coli* K-12. J Bacteriol, 120, 638–650.

30. Ueguchi, C., Suzuki, T., Yoshida, T., Tanaka, K. and Mizuno, T. (1996) Systematic mutational analysis revealing the functional domain organization of *Escherichia coli* nucleoid protein H-NS. J Mol Biol, 263, 149–162.

31. Kikhney, A.G. and Svergun, D.I. (2015) A practical guide to small angle X-ray scattering (SAXS) of flexible and intrinsically disordered proteins. FEBS Lett, 589, 2570–2577.

32. Jerabek-Willemsen, M., Andr√©, T., Wanner, R., Roth, H.M., Duhr, S., Baaske, P. and Breitsprecher, D. (2014) MicroScale Thermophoresis: Interaction analysis and beyond. Journal of Molecular Structure, 1077, 101–113.

33. Gordon, B.R., Li, Y., Cote, A., Weirauch, M.T., Ding, P., Hughes, T.R., Navarre, W.W., Xia, B. and Liu, J. (2011) Structural basis for recognition of AT-rich DNA by unrelated xenogeneic silencing proteins. Proc Natl Acad Sci U S A, 108, 10690–10695.

34. Nunn, C.M., Garman, E. and Neidle, S. (1997) Crystal structure of the DNA decamer d(CGCAATTGCG) complexed with the minor groove binding drug netropsin. Biochemistry, 36, 4792–4799.

35. Leung, C., Bestembayeva, A., Thorogate, R., Stinson, J., Pyne, A., Marcovich, C., Yang, J., Drechsler, U., Despont, M., Jankowski, T. et al. (2012) Atomic force microscopy with nanoscale cantilevers resolves different structural conformations of the DNA double helix. Nano Lett, 12, 3846–3850.

36. Cordeiro, T.N., Schmidt, H., Madrid, C., Juarez, A., Bernado, P., Griesinger, C., Garcia, J. and Pons, M. (2011) Indirect DNA readout by an H-NS related protein: structure of the DNA complex of the C-terminal domain of Ler. PLoS Pathog, 7, e1002380.

37. Ueguchi, C., Seto, C., Suzuki, T. and Mizuno, T. (1997) Clarification of the dimerization domain and its functional significance for the *Escherichia coli* nucleoid protein H-NS. J Mol Biol, 274, 145–151.

38. Ali, S.S., Whitney, J.C., Stevenson, J., Robinson, H., Howell, P.L. and Navarre, W.W. (2013) Structural insights into the regulation of foreign genes in *Salmonella* by the Hha/H-NS complex. J Biol Chem, 288, 13356–13369.

39. Madrid, C., Nieto, J.M. and Juarez, A. (2002) Role of the Hha/YmoA family of proteins in the thermoregulation of the expression of virulence factors. Int J Med Microbiol, 291, 425–432.

40. Paytubi, S., Madrid, C., Forns, N., Nieto, J.M., Balsalobre, C., Uhlin, B.E. and Juarez, A. (2004) YdgT, the Hha paralogue in *Escherichia coli*, forms heteromeric complexes with H-NS and StpA. Mol Microbiol, 54, 251–263.

41. Banos, R.C., Vivero, A., Aznar, S., Garcia, J., Pons, M., Madrid, C. and Juarez, A. (2009) Differential regulation of horizontally acquired and core genome genes by the bacterial modulator H-NS. PLoS Genet, 5, e1000513.

42. Vivero, A., Banos, R.C., Mariscotti, J.F., Oliveros, J.C., Garcia-del Portillo, F., Juarez, A. and Madrid, C. (2008) Modulation of horizontally acquired genes by the Hha-YdgT proteins in *Salmonella enterica* serovar Typhimurium. J Bacteriol, 190, 1152–1156.

43. Neves, M.A., Yeager, M. and Abagyan, R. (2012) Unusual arginine formations in protein function and assembly: rings, strings, and stacks. J Phys Chem B, 116, 7006–7013.

44. Lee, D., Lee, J. and Seok, C. (2013) What stabilizes close arginine pairing in proteins? Phys Chem Chem Phys, 15, 5844–5853.

45. Garcia-Perez, M., Pinto, M. and Subirana, J.A. (2003) Nonsequence-specific arginine interactions in the nucleosome core particle. Biopolymers, 69, 432–439.

46. Magalhaes, A., Maigret, B., Hoflack, J., Gomes, J.N. and Scheraga, H.A. (1994) Contribution of unusual arginine-arginine short-range interactions to stabilization and recognition in proteins. J Protein Chem, 13, 195–215.

47. Yamanaka, Y., Winardhi, R.S., Yamauchi, E., Nishiyama, S.I., Sowa, Y., Yan, J., Kawagishi, I., Ishihama, A. and Yamamoto, K. (2018) Dimerization site 2 of the bacterial DNA-binding protein H-NS is required for gene silencing and stiffened nucleoprotein filament formation. J Biol Chem, 293, 9496–9505.

48. Sievers, F., Wilm, A., Dineen, D., Gibson, T.J., Karplus, K., Li, W., Lopez, R., McWilliam, H., Remmert, M., Soding, J. et al. (2011) Fast, scalable generation of high-quality protein multiple sequence alignments using Clustal Omega. Mol Syst Biol, 7, 539.

49. Bertin, P., Benhabiles, N., Krin, E., Laurent-Winter, C., Tendeng, C., Turlin, E., Thomas, A., Danchin, A. and Brasseur, R. (1999) The structural and functional organization of H-NS-like proteins is evolutionarily conserved in gram-negative bacteria. Mol Microbiol, 31, 319–329.

50. Grabarek, Z. and Gergely, J. (1990) Zero-length crosslinking procedure with the use of active esters. Anal Biochem, 185, 131–135.

51. Schindelin, J., Arganda-Carreras, I., Frise, E., Kaynig, V., Longair, M., Pietzsch, T., Preibisch, S., Rueden, C., Saalfeld, S., Schmid, B. et al. (2012) Fiji: an open-source platform for biological-image analysis. Nat Methods, 9, 676-682.

