## Supporting material for "Deconstructing the structural conservation of distantly related bacterial nucleoid-associated proteins using functional chimeras"

A

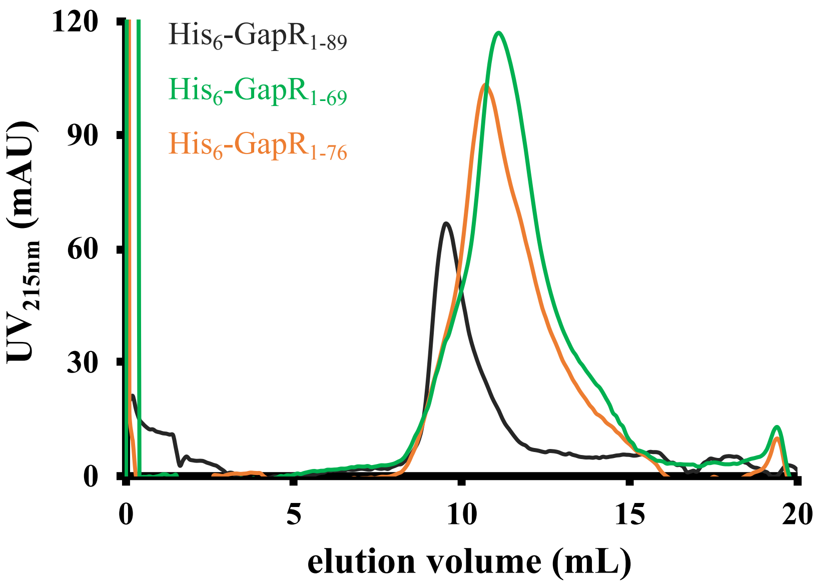


| **Protein** | **Stoichiometry** |
|---|---|
| His$_6$-GapR$_{1-89}$ | 4.73 x |
| His$_6$-GapR$_{1-69}$ | 2.92 x |
| His$_6$-GapR$_{1-76}$ | 3.13 x |

B

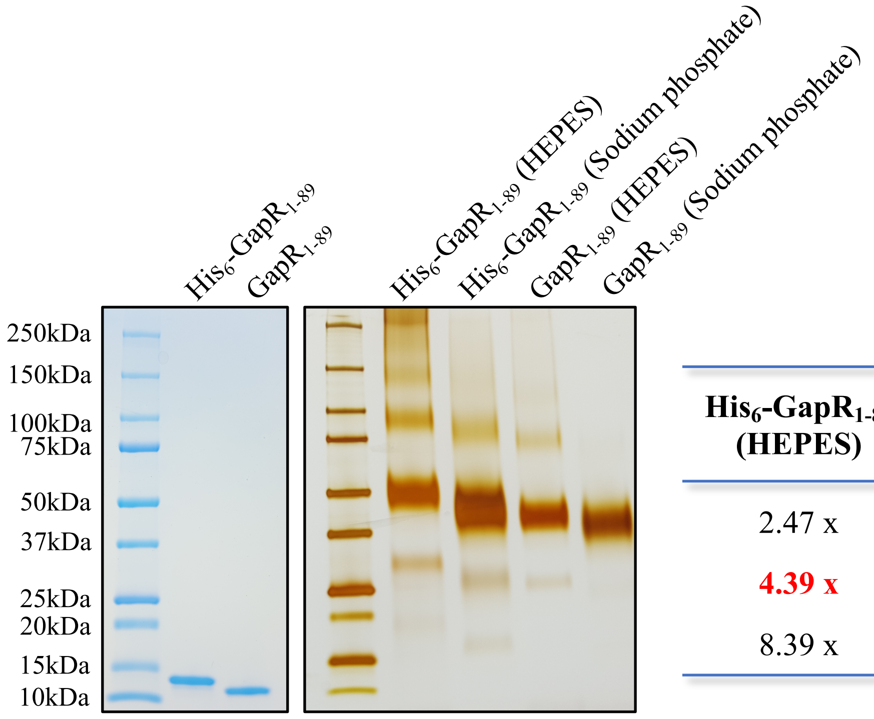


| **His$_6$-GapR$_{1-89}$ (HEPES)** | **His$_6$-GapR$_{1-89}$ (Sodium phosphate)** | **GapR$_{1-89}$ (HEPES)** | **GapR$_{1-89}$ (Sodium phosphate)** | |
|---|---|---|---|---|
| 2.47 x | 2.13 x | 2.44 | | dimer |
| **4.39 x** | **3.81 x** | **4.28** | **4.10** | tetramer |
| 8.39 x | 7.47 x | 8.19 | | octamer |

**Figure S1.** GapR is a tetramer in solution. **A)** Determination of the oligomeric state of GapR proteins. Affinity purified proteins were separated from co-purified DNA and analyzed by size exclusion chromatography using column Superdex 75 10/300 GL in the presence of 1M NaCl and 1 mM EDTA. The protein samples were analyzed at the following concentrations: His$_6$-GapR$_{1-89}$ at 50 µM (monomer), His$_6$-GapR$_{1-69}$ at 125 µM (monomer) and His$_6$-GapR$_{1-76}$ at 100 µM (monomer). The apparent molecular weight of each elution peak was estimated and divided by the molecular weight calculated for the monomeric state of the corresponding protein. **B)** Neither the His-tag nor sodium phosphate alters the oligomeric state of GapR$_{1-89}$. Tagged and untagged GapR$_{1-89}$ were dialyzed against either 20 mM HEPES, 150 mM NaCl, 10% glycerol or 50 mM sodium phosphate, 150 mM NaCl, 10% glycerol and assayed in crosslinking reactions. Proteins at 50 µM (monomer) were treated with 400 mM EDC + 100 mM NHS for 2 h at room temperature, the reaction products were resolved by SDS-PAGE, and the gel was silver stained. As a control, proteins incubated for 2 h at room temperature in the absence of the crosslinking agents were resolved by SDS-PAGE, and the gel was stained with Coomassie blue. The apparent molecular weight of each band was estimated and divided by the molecular weight calculated for the monomeric state of the corresponding protein. Values denoted in red refer to the main bands detected in the gel. The figure shows that phosphate causes both tagged and untagged GapR$_{1-89}$ to migrate slightly faster in SDS-PAGE compared to the proteins in HEPES buffer.

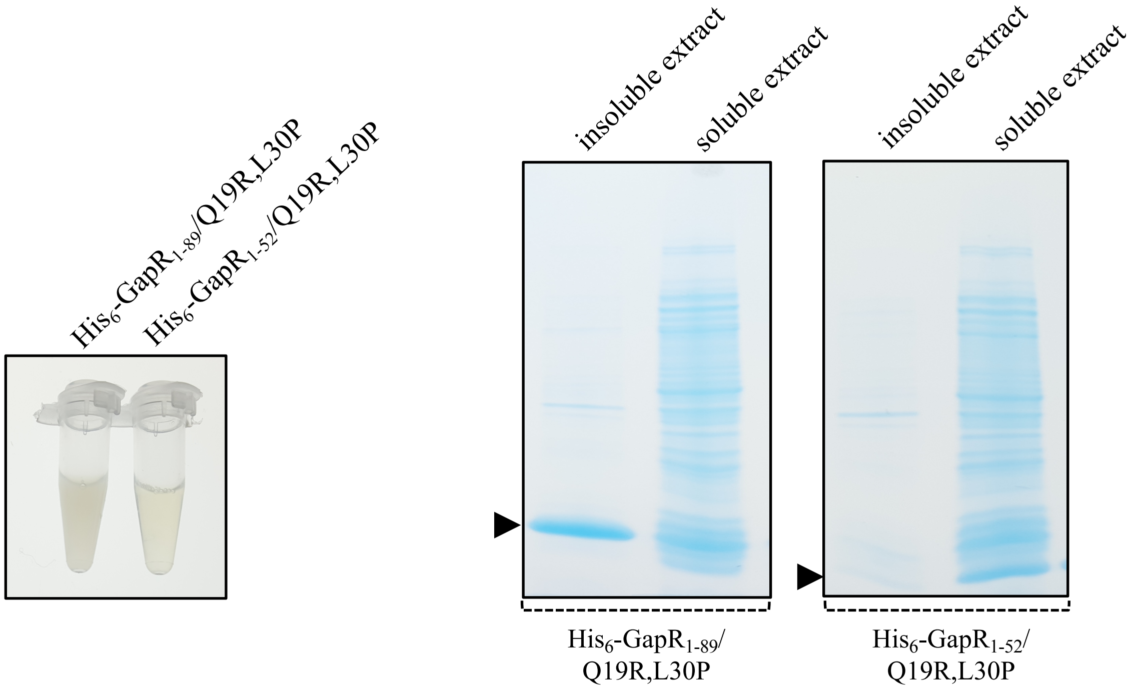


**Figure S2.** Deletion of the C-terminal region renders GapR/Q19R,L30P more soluble. Evaluation of the solubility of GapR proteins by SDS-PAGE. On the left, suspensions of *E. coli* expressing either $His_6$-$GapR_{1\text{-}89}$/Q19R,L30P or $His_6$-$GapR_{1\text{-}52}$/Q19R,L30P were sonicated and imaged. On the right, soluble and insoluble fractions prepared after cell lysis were resolved by SDS-PAGE, and the gel was stained with Coomassie blue. The proteins are indicated by arrowheads.

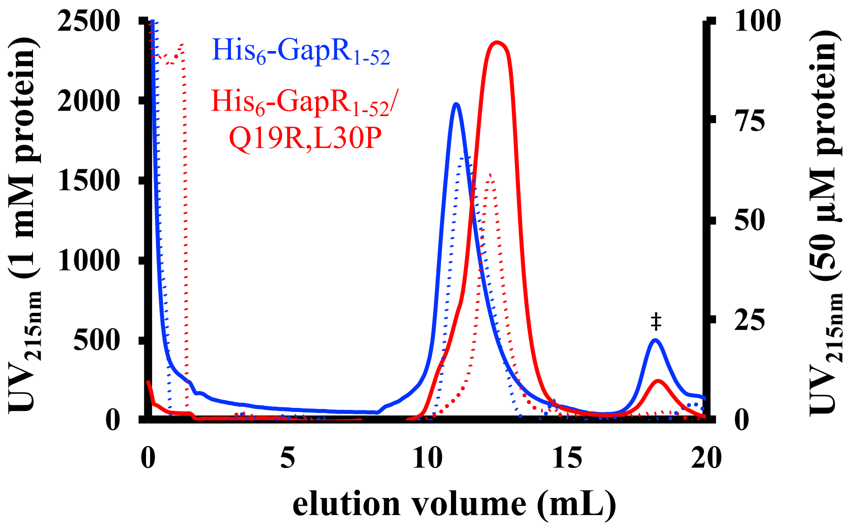


| Protein (concentration) | Stoichiometry |
|---|---|
| His$_6$-GapR$_{1\text{-}52}$ (50 µM) | 3.31 x |
| His$_6$-GapR$_{1\text{-}52}$ (1 mM) | 3.84 x |
| His$_6$-GapR$_{1\text{-}52}$/Q19R,L30P (50 µM) | 2.41 x |
| His$_6$-GapR$_{1\text{-}52}$/Q19R,L30P (1 mM) | 2.25 x |

**Figure S3.** GapR protein truncations corresponding to the coiled-coil motif in the N-terminus were eluted with apparent molecular weights higher than those expected. Affinity purified His-tagged proteins at the concentrations of 50 µM (monomer) (solid lines) and 1 mM (monomer) (dotted lines) were analyzed by size exclusion chromatography using column Superdex 75 10/300 GL in the presence of 150 mM NaCl. No protein was detected on the peak denoted by ‡. The apparent molecular weight of each elution peak was estimated and divided by the molecular weight calculated for the monomeric state of the corresponding protein.

A

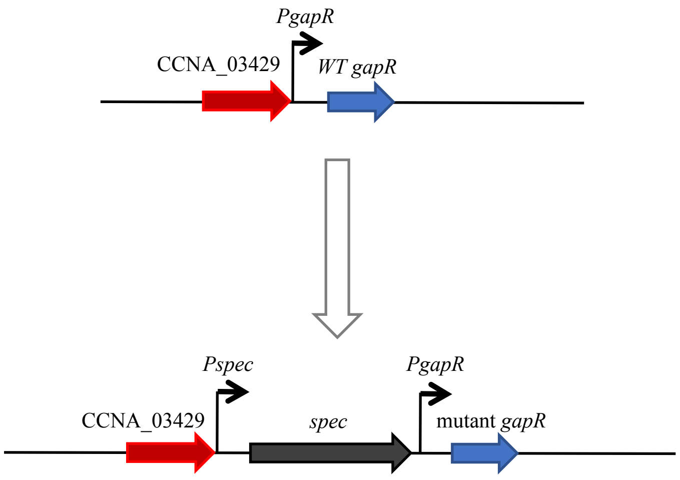


| Residue(s) | Time after plating | # analyzed | # replaced |
|---|---|---|---|
| K59 | 3 days | 40 | 26 |
| R65A,K66A | 3 days | 40 | 0 |
| | 7 days | 40 | 40 |
| Q19R,L30P | 3 days | 40 | 0 |
| | 5 days | 40 | 40 |

B

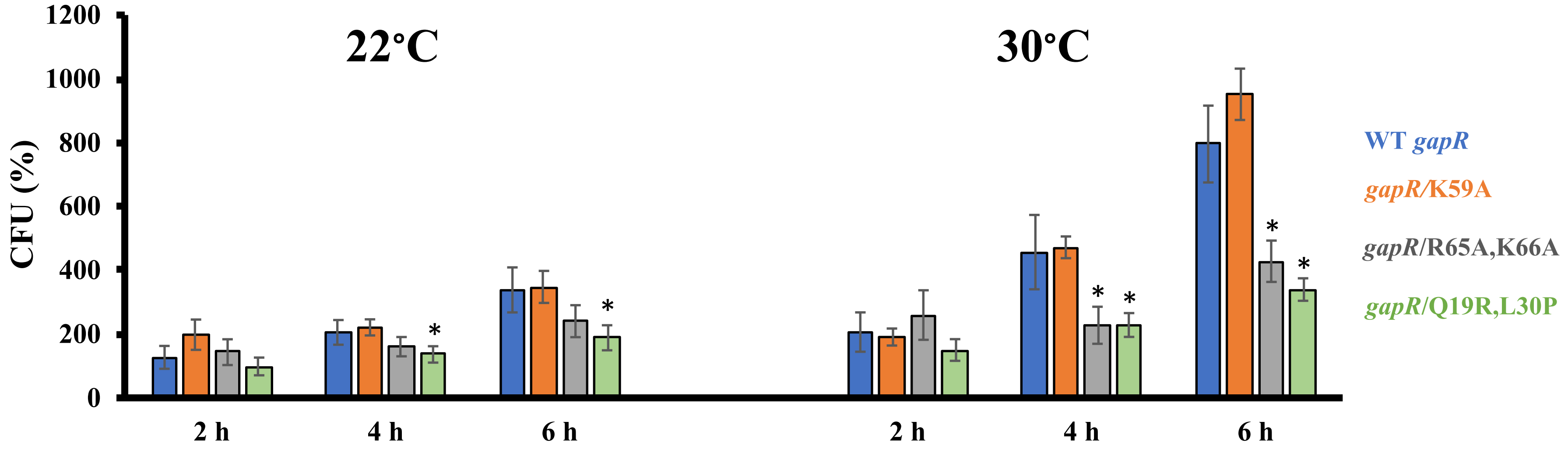


C

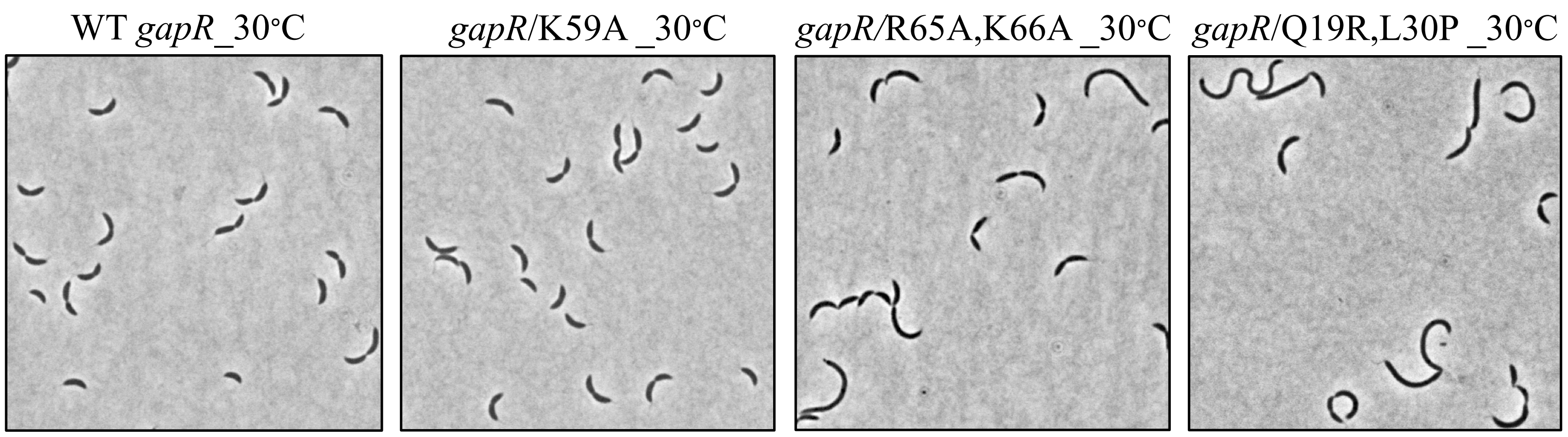


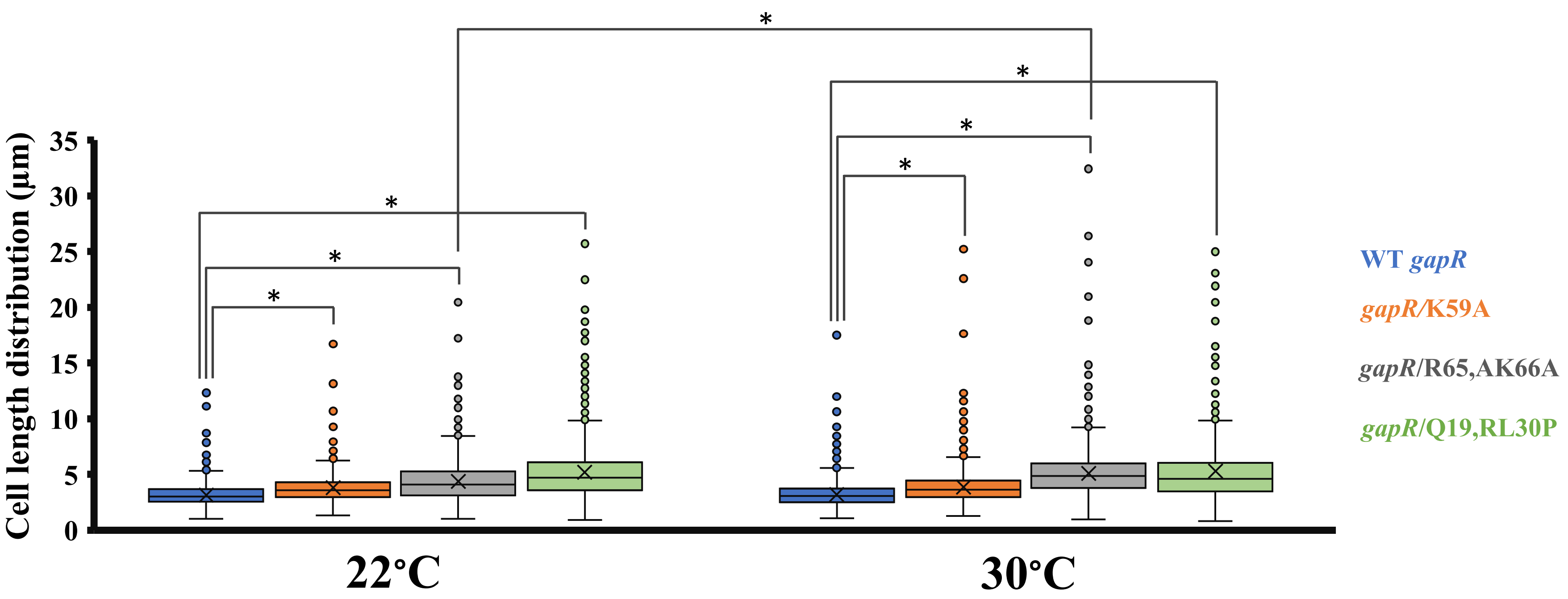


**Figure S4.** Legend on next page

**Figure S4.** Mutations selectively disrupting oligomerization or DNA binding activity of GapR reduce cell growth and increase cell length. **A)** Left panel: schematic of the strategy used to replace wild-type *gapR* for a mutant copy of the gene. Right panel: screening for *gapR* mutants. Colonies at the indicated time point after plating in the selection medium were screened. For each strain, the number of colonies analyzed and the number of mutant colonies identified are represented. **B)** Growth analysis of mutant *gapR* strains. Saturated overnight cultures grown at 22°C were diluted to $OD_{600nm}$ of 0.1 and incubated at both 22 and 30°C. At the indicated time points, aliquots were taken for counting colony forming units (CFU). Values are the percentage of colony forming units at the indicated time points (t = 2, 4 and 6 h) relative to that determined for the freshly diluted cultures (t = 0). Results are mean of three independent biological experiments, and bars indicate standard deviations. **C)** Cell morphology analysis of mutant *gapR* strains. Exponentially growing cells ($OD_{600nm}$ of 0.3-0.5) at both 22 and 30°C were imaged by phase-contrast microscopy. The length of individual cells from each strain was determined by the MicrobeJ tool (1), and data analysis was performed using the R statistical program (Table S3). The numbers of individual cells used for analysis were as follows: WT *gapR*_22°C = 3343, *gapR*/K59A_22°C = 1226, *gapR*/R65A,K66A_22°C = 1585, *gapR*/Q19R,L30P_22°C = 1645, WT *gapR*_30°C = 2378, *gapR*/K59A_30°C = 1235, *gapR*/R65A,K66A_30°C = 1477, *gapR*/Q19R,L30P_30°C = 690. Representative images of each strain grown at 30°C are shown. For both panels, asterisk represents statistically different values according to unpaired Student's t-Test ($p<0.001$).

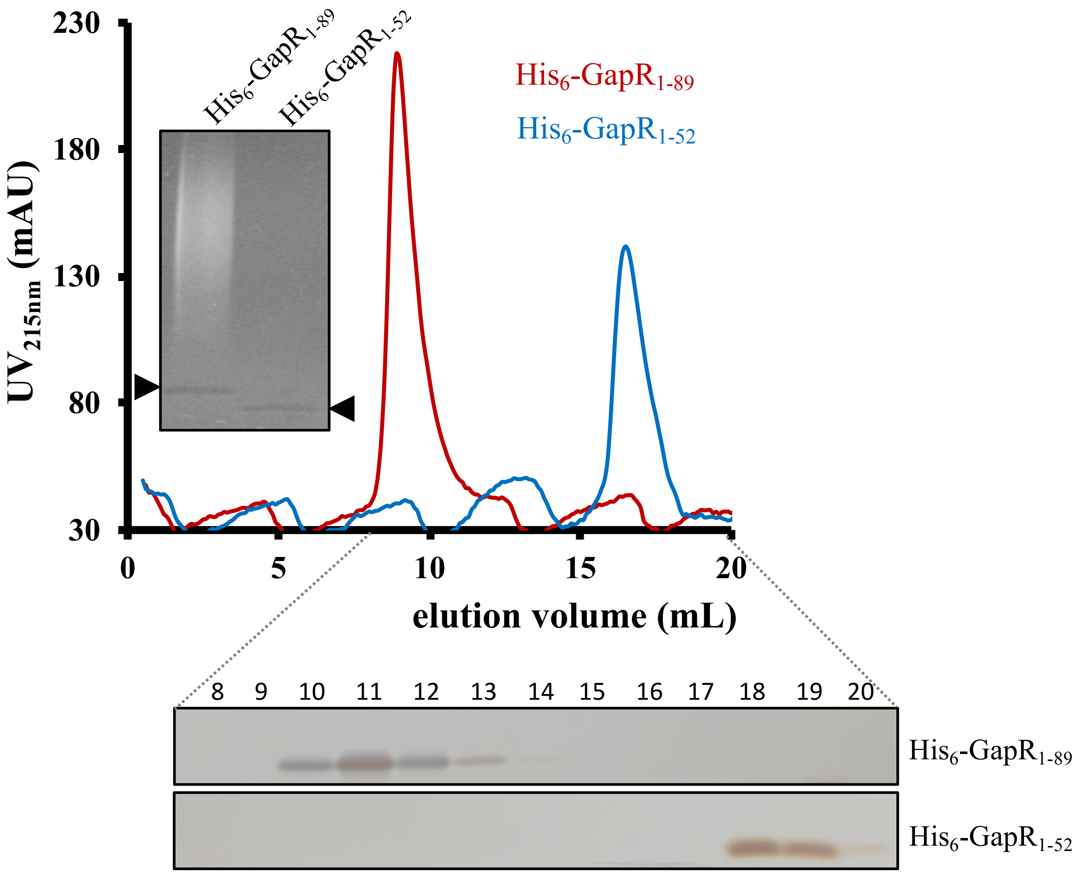


**Figure S5.** The C-terminal region of GapR is essential for the DNA-binding activity of the protein. His$_6$-GapR$_{1\text{-}89}$ and His$_6$-GapR$_{1\text{-}52}$ purified without a nuclease treatment were analyzed by size exclusion chromatography using column Superdex 200 10/300 GL in the presence of 150 mM NaCl. 50 µM protein (monomer) was used for each run. Fractions 8-20 were resolved by SDS-PAGE, and the gels were silver stained. Inset: SDS-PAGE of the affinity-purified proteins. The gel was stained with Coomassie blue and then with ethidium bromide. Arrows indicate proteins, and the smear corresponds to co-purified DNA, which is only visible in the His$_6$-GapR$_{1\text{-}89}$ sample.

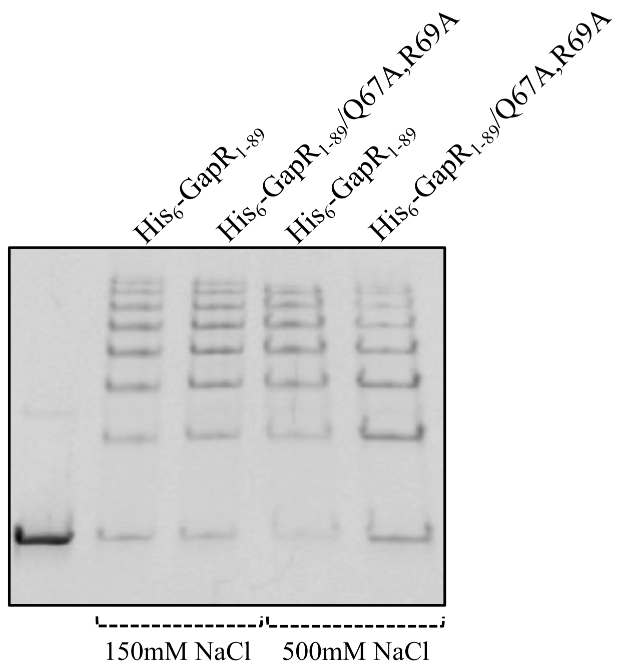


**Figure S6.** Substitution of non-conserved, polar residues at the C-terminal region of GapR only effects the DNA-binding activity of the protein under high ionic strength condition. The DNA-binding affinity of the proteins was monitored by electrophoretic mobility shift assays. 2.5 μM protein (monomer) was incubated with 0.1 μM 320 bp $P_{pilA}$ DNA for 30 min at room temperature in the presence of either 150 or 500 mM NaCl, the reaction products were resolved by PAGE under native conditions, and the gel was stained with ethidium bromide. A control with no protein included is also shown.

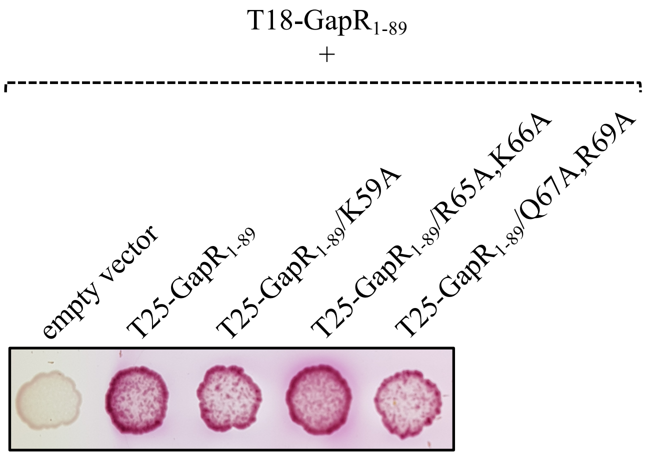


**Figure S7.** Mutations in the DNA-binding motif do not affect GapR oligomerization. The ability of mutant GapR proteins to interact with wild-type GapR was analyzed by bacterial two-hybrid assays. *E coli* BTH101 cells expressing the T18 domain of adenylate cyclase N-terminally fused to wild-type full-length GapR and the T25 domain of the same enzyme fused to the N-terminus of different mutant full-length GapR proteins were grown to exponential phase ($OD_{600nm}$ of 0.5), and 3 μL from each culture were spotted in 1% maltose-containing MacConkey plates. The plates were imaged after 2 days at 30°C.

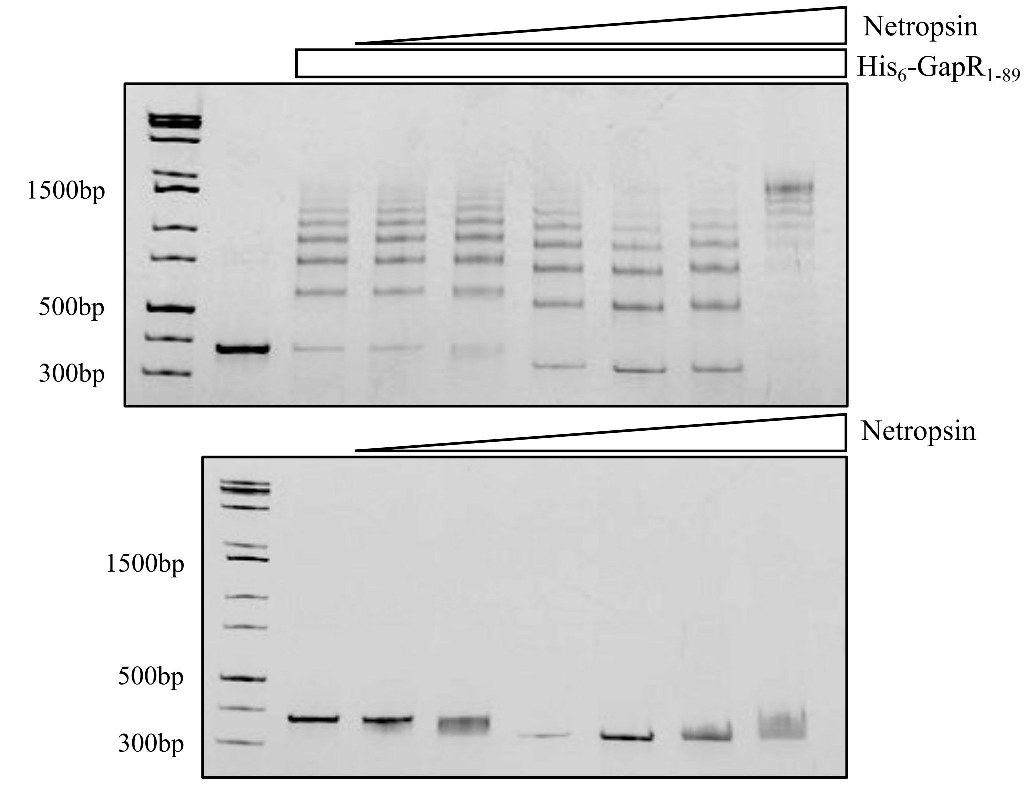


**Figure S8.** GapR does not compete with Netropsin for binding DNA. The DNA-binding affinity of His$_6$-GapR$_{1\text{-}89}$ was monitored by electrophoretic mobility shift assays. 0.1 µM 320 bp P$_{pilA}$ DNA was treated with different concentrations of Netropsin (10 nM to 1 mM) for 30 min at room temperature in the presence of 150 mM NaCl. 2.5 µM His$_6$-GapR$_{1\text{-}89}$ (monomer) was added, and the resulting mixtures were incubated for an additional 30 min at room temperature. Controls with no His$_6$-GapR$_{1\text{-}89}$ included were performed to evaluate the binding of Netropsin to DNA. The reaction products were resolved by PAGE under native conditions, and the gels were stained with ethidium bromide.

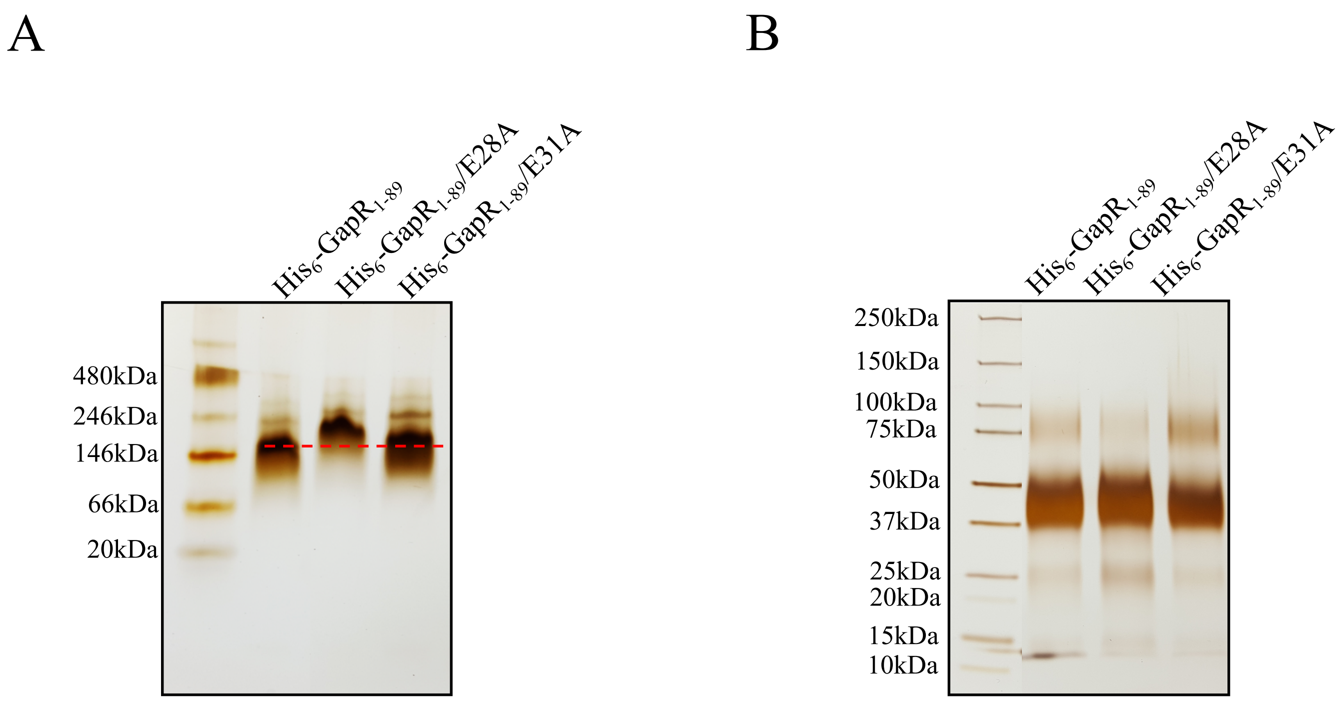


**Figure S9.** The E28A mutation alters the migration of GapR under native PAGE but not affects the oligomeric state of the protein. Affinity purified proteins were separated from co-purified DNA, dialyzed against buffer containing 150 mM NaCl and resolved by native PAGE **(A)** or used in a crosslinking reaction **(B)**. For the native PAGE, proteins at 10 µM (monomer) were used. For the crosslinking reaction, proteins at 50 µM (monomer) were treated with 400 mM EDC + 100 mM NHS for 2 h at room temperature, and the reaction products were resolved by SDS-PAGE. The gels were silver stained. His$_6$-GapR/E31A was used as a control. The E31 residue is solvent exposed, so no structural change is expected upon replacing it to alanine. Therefore, this residue is appropriate to determine the impact of losing one negative charge on the electrophoretic migration of GapR.

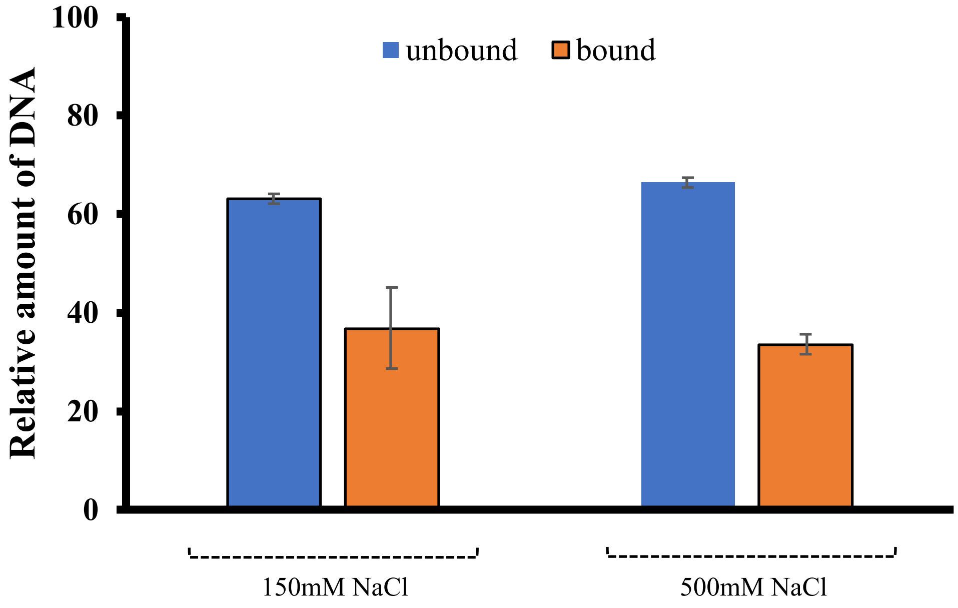


**Figure S10.** GapR binds DNA that is not overtwisted due to its high AT content. Electrophoretic mobility shift assay performed using 10 µM protein (monomer) and 1 µM 12-mer DNA fragment $(dCG)_6$. Protein and DNA were incubated for 30 min at room temperature, the reaction products were resolved by PAGE under native conditions, and the gels were stained with ethidium bromide. Controls with no protein added were included. The relative amount of DNA in both unbound and bound states were quantified using Image J (2).

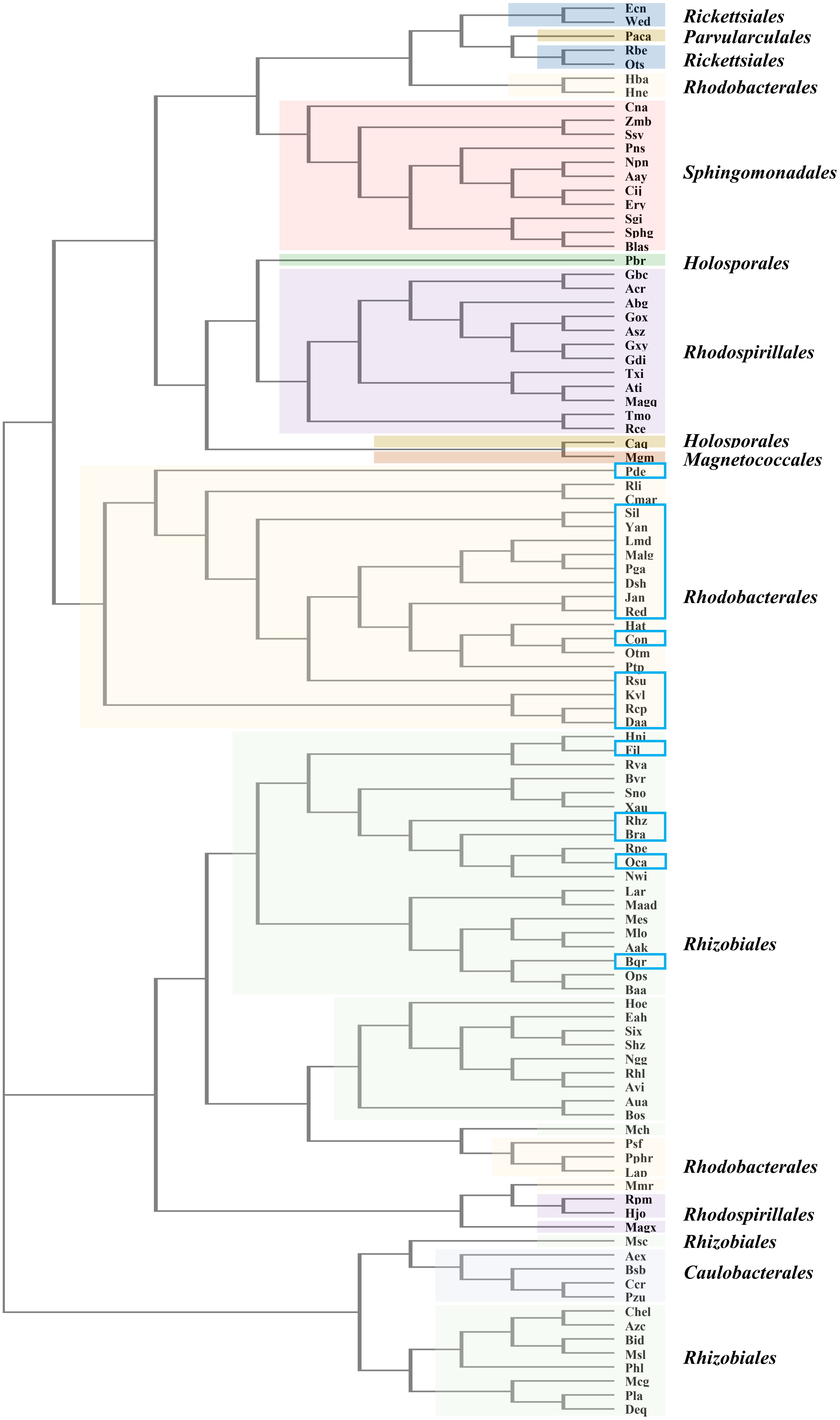


**Figure S11.** Legend on next page

**Figure S11.** GapR but not H-NS is widespread in the α subdivision of Proteobacteria. Distribution of H-NS-like proteins among selected GapR-containing alphaproteobacterial species. H-NS orthologs were identified by mining the Kyoto Encyclopedia of Genes and Genomes (KEGG) (3) database for amino acid sequences possessing the Pfam domain Histone_HNS and displaying significant identity (threshold for Smith-Waterman score of 200) to the trans-acting regulatory protein HvrA from *Rhodobacter capsulatus* SB 1003, a H-NS-like protein (4,5). The occurrence of H-NS-like proteins is represented by blue rectangles in a neighbor-joining phylogenetic tree constructed using the Clustal Omega tool (6). Species abbreviations: *Caulobacter crescentus* NA1000 (Ccr; YP_002518801) *Phenylobacterium zucineum* HLK1 (Pzu; ACG79526), *Brevundimonas subvibrioides* ATCC 15264 (Bsb; ADL02288), *Asticcacaulis excentricus* CB 48 (Aex; ADU13212), *Parvibaculum lavamentivorans* DS-1 (Pla; ABS63648), *Mesorhizobium japonicum* MAFF 303099 (Mlo; BAB50739), *Devosia* sp. H5989 (Deq; AKR57895), *Aureimonas* sp. AU20 (Aua; ALN74248), *Bosea* sp. RAC05 (Bos; AOG04397), *Methylobacterium extorquens* CM4 (Mch; ACK83815), *Chelativorans* sp. BNC1 (Mes; ABG64738), *Oligotropha carboxidovorans* OM5 (Oca; ACI92221), *Pelagibacterium halotolerans* B2 (Phl; AEQ52522), *Aminobacter aminovorans* (Aak; AMS40393), *Bradyrhizobium* sp. ORS 278 (Bra; CAL75095), *Starkeya novella* DSM 506 (Sno; ADH90386), *Blastochloris viridis* (Bvr; ALK07853), *Martelella* sp. AD-3 (Maad; AMM85466), *Methylocystis* sp. SC2 (Msc; CCJ06210), *Beijerinckia indica* subsp. *indica* ATCC 9039 (Bid; ACB96787), *Rhizobium favelukesii* (Rhl; CDM59164), *Sinorhizobium* sp. RAC02 (Six; AOF92024), *Methyloceanibacter caenitepidi* (Mcg; BAQ16066), *Nitrobacter winogradskyi* Nb-255 (Nwi; ABA04251), *Shinella* sp. HZN7 (Shz; ANH05253), *Rhodopseudomonas palustris* BisA53 (Rpe; ABJ08033), *Agrobacterium vitis* S4 (Avi; ACM37657), *Ochrobactrum pseudogrignonense* (Ops; ANG96594), *Ensifer adhaerens* (Eah; ANK73839), *Hoeflea* sp. IMCC20628 (Hoe; AKH99110), *Neorhizobium galegae* bv. orientalis str. HAMBI 540 (Ngg; CDN49226), *Xanthobacter autotrophicus* Py2 (Xau; ABS67310), *Rhodomicrobium vannielii* ATCC 17100 (Rva; ADP70897), *Candidatus Liberibacter americanus* str. Sao Paulo (Lar; AHA28099), *Chelatococcus* sp. CO-6 (Chel; ALA16615), *Bartonella quintana* RM-11 (Bqr; AFR26807), *Azorhizobium caulinodans* ORS 571 (Azc; BAF86439), *Hyphomicrobium nitrativorans* NL23 (Hni; AHB49456), *Methylocella silvestris* BL2 (Msl; ACK52583), *Rhodoplanes* sp. Z2-YC6860 (Rhz; AMN40225), *Candidatus Filomicrobium marinum* (Fil; CFX29357), *Brucella abortus* A13334 (Baa; AEW17098), *Hirschia baltica* ATCC 49814 (Hba; ACT60610), *Pannonibacter phragmitetus* (Pphr; ALV27237), *Labrenzia* sp. CP4 (Lap; AMN51817), *Pseudovibrio* sp. FO-BEG1 (Psf; AEV39265), *Maricaulis maris* MCS10 (Mmr; ABI66821), *Hyphomonas neptunium* ATCC 15444 (Hne; ABI77018), *Defluviimonas alba* (Daa; AMY69002), *Ketogulonicigenium vulgare* WSH-001 (Kvl; YP_005795322), *Ruegeria pomeroyi* DSS-3 (Sil; AAV94337), *Yangia* sp. CCB-MM3 (Yan; ANT62436), *Roseobacter litoralis* Och 149 (Rli; AEI93600), *Confluentimicrobium* sp. EMB200-NS6 (Con; ALG90916), *Paracoccus denitrificans* PD1222 (Pde; ABL71804), *Marinovum algicola* DG 898 (Malg; AKO97568), *Celeribacter marinus* (Cmar; ALI56317), *Octadecabacter temperatus* (Otm; AKS45751), *Phaeobacter inhibens* DSM 17395 (Pga; AFO91549), *Rhodovulum sulfidophilum* (Rsu; BAQ70137), *Planktomarina temperata* RCA23 (Ptp; AII85964), *Halocynthiibacter arcticus* (Hat; AML53785), *Rhodobacter capsulatus* SB 1003 (Rcp; ADE85717), *Dinoroseobacter shibae* DFL 12 = DSM 16493 (Dsh; ABV93161), *Leisingera methylohalidivorans* DSM 14336 (Lmd; AHD00583), *Jannaschia* sp. CCS1 (Jan; ABD55552), *Roseibacterium elongatum* DSM 19469 (Red; AHM05525), *Sphingorhabdus* sp. M41 (Sphg; AMO70626), *Sphingobium* sp. SYK-6 (Ssy; BAK65186), *Croceicoccus naphthovorans* (Cna; AKM11472), *Sphingopyxis granuli* (Sgi; AMG73626), *Zymomonas mobilis* subsp. *mobilis* ATCC 29191 (Zmb; AFN56942), *Blastomonas* sp. RAC04 (Blas; AOG02067), *Novosphingobium pentaromativorans* US6-1 (Npn; AIT80101), *Altererythrobacter atlanticus* (Aay; AKH42983), *Citromicrobium* sp. JL477 (Cij; ALG59616), *Erythrobacter atlanticus* (Ery; AKQ40906), *Porphyrobacter neustonensis* (Pns; ANK12064), *Azospirillum thiophilum* (Ati; ALG71691), *Rhodospirillum centenum* SW (Rce; ACI99528), *Magnetospirillum* sp. XM-1 (Magx; CUW40179), *Magnetospira* sp. QH-2 (Magq; CCQ75212), *Granulibacter bethesdensis* (Gbc; AHJ62133), *Thalassospira xiamenensis* M-5 = DSM 17429 (Txi; AJD51518), *Komagataeibacter medellinensis* NBRC 3288 (Gxy; BAK83223), *Acidiphilium cryptum* JF-5 (Acr; ABQ29408), *Gluconobacter oxydans* 621H (Gox; AAW59937), *Acetobacter senegalensis* (Asz; CEF40008), *Gluconacetobacter diazotrophicus* PA1 5 (Gdi; CAP57119), *Pararhodospirillum photometricum* DSM 122 (Rpm; CCG09558), *Haematospirillum jordaniae* (Hjo; AMW35670), *Tistrella mobilis* KA081020-065 (Tmo; AFK56936), *Asaia bogorensis* NBRC 16594 (Abg; BAT19319), *Orientia tsutsugamushi* str. Boryong (Ots; CAM79926), *Rickettsia bellii* RML369-C (Rbe; ABE05266), *Wolbachia endosymbiont of Drosophila simulans wNo* (Wed; AGJ98833), *Ehrlichia canis* str. Jake (Ecn; AAZ68499), *Parvularcula bermudensis* HTCC2503 (Pbr; ADM10025), *Candidatus Paracaedimonas acanthamoebae* (Caq; AIL12384), *Candidatus Paracaedibacter acanthamoebae* (Paca; AIK97255), *Magnetococcus marinus* MC-1 (Mgm; ABK43434).

**Table S1.** Strains

| Strain | Description | Source |
|---|---|---|
| ***E.coli*** | | |
| NEB Turbo | *F' proA$^{+}$B$^{+}$ lacI$^{q}$ ΔlacZM15/fhuA2 Δ(lac-proAB) glnV galK16 galE15 R(zgb-210::Tn10)Tet$^{S}$ endA1 thi-1 Δ(hsdS-mcrB)5*; for cloning | New England Biolabs |
| BL21 (DE3) | *fhuA2 [lon] ompT gal (λ DE3) [dcm] ΔhsdS λ DE3 = λ sBamHIo ΔEcoRI-B int::(lacI::PlacUV5::T7 gene1) i21 Δnin5; for protein expression* | New England Biolabs |
| Δ*hns* | MG1655 *hns*::*tetRA*; for *bgl* assay | (7) |
| BTH101 | *F$^{-}$ cya-99 araD139 galE15 galK16 rpsL1 hsdR2 mcrA1 mcrB1*; for bacterial two-hybrid assay | Euromedex |
| ***C. crescentus*** | | |
| NA1000 | Holdfast mutant derivative of wild-type CB15; for phenotypic analyses | (8) |
| *gapR*/Q19R,L30P | NA1000 *gapR*/Q19R,L30P, Spec$^{R}$; for phenotypic analyses | This work |
| *gapR*/K59A | NA1000 *gapR*/K59A, Spec$^{R}$; for phenotypic analyses | This work |
| *gapR*/R65A,K66A | NA1000 *gapR*/R65A,K66A, Spec$^{R}$; for phenotypic analyses | This work |

**Table S2.** Plasmids

| Plasmid | Description | Source |
|---|---|---|
| pBAD33 | Replicating vector for arabinose-inducible expression; Cm$^{R}$ | (9) |
| pKT25 | Replicating vector for IPTG-inducible expression of CyaA$^{T25}$ fused to the N terminus of a protein of interest; Kan$^{R}$ | Euromedex |
| pUT18C | Replicating vector for PTG-inducible expression of CyaA$^{T18}$ fused to the N terminus of a protein of interest; Amp$^{R}$ | Euromedex |
| pET28b | Replicating vector for IPTG-inducible expression of His$_{6}$ fused to the N terminus of a protein of interest; Kan$^{R}$ | Novagen |
| pNPTS138 | Suicide vector for two-step recombination; Kan$^{R}$, Suc$^{S}$ | (10) |
| pBAD33*gapR*1-89 | For arabinose-inducible expression of GapR$_{1-89}$ | (11) |
| pBAD33*hns*1-137 | For arabinose-inducible expression of H-NS$_{1-137}$ | (11) |
| pBAD33*gapR*1-49-*hns*85-137 | For arabinose-inducible expression of GapR$_{1-49}$-H-NS$_{85-137}$ | This work |
| pBAD33*gapR*1-47-*hns*51-137 | For arabinose-inducible expression of GapR$_{1-47}$-H-NS$_{51-137}$ | This work |
| pBAD33*hns*1-22-*gapR*20-49-*hns*85-137 | For arabinose-inducible expression of H-NS$_{1-22}$-GapR$_{20-49}$-H-NS$_{85-137}$ | This work |
| pBAD33*hns*1-22-*gapR*20-47-*hns*51-137 | For arabinose-inducible expression of H-NS$_{1-22}$-GapR$_{20-47}$-H-NS$_{51-137}$ | This work |

| | | |
|---|---|---|
| pBAD33*hns*1-84 | For arabinose-inducible expression of H-$NS_{1-84}$ | This work |
| pBAD33*hns*1-84-*gapR*50-89 | For arabinose-inducible expression of H-$NS_{1-84}$-$GapR_{50-89}$ | This work |
| pBAD33*hns*1-110-*gapR*50-89 | For arabinose-inducible expression of H-$NS_{1-110}$-$GapR_{50-89}$ | This work |
| pBAD33*hns*Δ23-84 | For arabinose-inducible expression of H-NS with an internal deletion corresponding to residues 23-84 | This work |
| pBAD33*hns*Δ23-50 | For arabinose-inducible expression of H-NS with an internal deletion corresponding to residues 23-50 | This work |
| pBAD33*hns*Δ51-84 | For arabinose-inducible expression of H-NS with an internal deletion corresponding to residues 51-84 | This work |
| pUT18C*gapR*1-89 | For IPTG-inducible expression of $CyaA^{T18}$-$GapR_{1-89}$ | (11) |
| pKT25*gapR*1-89 | For IPTG-inducible expression of $CyaA^{T25}$-$GapR_{1-89}$ | (11) |
| pKT25*gapR*1-89/I23N (M1) | For IPTG-inducible expression of $CyaA^{T25}$-$GapR_{1-89}$/I23N | This work |
| pKT25*gapR*1-89/Q19R,L30P (M2) | For IPTG-inducible expression of $CyaA^{T25}$-$GapR_{1-89}$/Q19R,L30P | This work |
| pKT25*gapR*1-89/L20R,I24N,K42E (M3) | For IPTG-inducible expression of $CyaA^{T25}$-$GapR_{1-89}$/L20R,I24N,K42E | This work |
| pKT25*gapR*1-89/K59A | For IPTG-inducible expression of $CyaA^{T25}$-$GapR_{1-89}$/K59A | This work |
| pKT25*gapR*1-89/R65A,K66A | For IPTG-inducible expression of $CyaA^{T25}$-$GapR_{1-89}$/R65A,K66A | This work |
| pKT25*gapR*1-89/Q67A,R69A | For IPTG-inducible expression of $CyaA^{T25}$-$GapR_{1-89}$/Q67A,R69A | This work |
| pKT25*gapR*1-52 | For IPTG-inducible expression of $CyaA^{T25}$-$GapR_{1-52}$ | This work |
| pKT25*gapR*1-52/I23N (M1) | For IPTG-inducible expression of $CyaA^{T25}$-$GapR_{1-52}$/I23N | This work |
| pKT25*gapR*1-52/Q19R,L30P (M2) | For IPTG-inducible expression of $CyaA^{T25}$-$GapR_{1-52}$/Q19R,L30P | This work |
| pKT25*gapR*1-52/L20R,I24N,K42E (M3) | For IPTG-inducible expression of $CyaA^{T25}$-$GapR_{1-52}$/L20R,I24N,K42E | This work |
| pET28b*gapR*1-89 | For IPTG-inducible expression of $His_6$-$GapR_{1-89}$ | This work |
| pET28b*gapR*1-89/Q19R,L30P | For IPTG-inducible expression of $His_6$-$GapR_{1-89}$/Q19R,L30P | This work |
| pET28b*gapR*1-89/K59A | For IPTG-inducible expression of $His_6$-$GapR_{1-89}$/K59A | This work |
| pET28b*gapR*1-89/R65A,K66A | For IPTG-inducible expression of $His_6$-$GapR_{1-89}$/R65A,K66A | This work |

| | | |
|---|---|---|
| pET28b*gapR*1-89/Q67A,R69A | For IPTG-inducible expression of $His_6$-$GapR_{1\text{-}89}$/Q67A,R69A | This work |
| pET28b*gapR*1-89/E28A | For IPTG-inducible expression of $His_6$-$GapR_{1\text{-}89}$/E28A | This work |
| pET28b*gapR*1-89/E31A | For IPTG-inducible expression of $His_6$-$GapR_{1\text{-}89}$/E31A | This work |
| pET28b*gapR*1-76 | For IPTG-inducible expression of $His_6$-$GapR_{1\text{-}76}$ | This work |
| pET28b*gapR*1-69 | For IPTG-inducible expression of $His_6$-$GapR_{1\text{-}69}$ | This work |
| pET28b*gapR*1-52 | For IPTG-inducible expression of $His_6$-$GapR_{1\text{-}52}$ | This work |
| pET28b*gapR*1-52/Q19R,L30P | For IPTG-inducible expression of $His_6$-$GapR_{1\text{-}52}$/Q19R,L30P | This work |
| pNPTS138*specgapR*/Q19R,L30P | For replacement of WT *gapR* with the mutant allele *gapR*/Q19R,L30P and insertion of the Ω cassette in NA1000 | This work |
| pNPTS138*specgapR*/K59A | For replacement of WT *gapR* with the mutant allele *gapR*/K59A and insertion of the Ω cassette in NA1000 | This work |
| pNPTS138*specgapR*/R65A,K66A | For replacement of WT *gapR* with the mutant allele *gapR*/R65A,K66A and insertion of the Ω cassette in NA1000 | This work |

**Table S3.** Statistical analysis of the data obtained by measuring the cell length of individual cells.

**Mean length (µm) of *C. crescentus* cells expressing different GapR proteins**

| **Temperature (°C)** | **WT *gapR*** | ***gapR*/K59A** | ***gapR*/R65A,K66A** | ***gapR*/Q19R,L30P** |
|---|---|---|---|---|
| 22°C | 3.148 | 3.786 | 4.355 | 5.177 |
| 30°C | 3.216 | 3.872 | 5.091 | 5.282 |

**Comparison of the cell length distribution of different strains at the same temperature**

| | **Temperature (°C)** | ***gapR*/K59A x WT** | ***gapR*/R65A,K66A x WT** | ***gapR*/Q19R,L30P x WT** |
|---|---|---|---|---|
| mean length difference (µm) (IC - 95%) | 22°C | 0.639 (0.564 to 0.713) | 1.207 (1.119 to 1.295) | 2.029 (1.897 to 2.161) |
| | 30°C | 0.656 (0.563 to 0.749) | 1.875 (1.761 to 1.989) | 2.066 (1.828 to 2.304) |

**Comparison of the cell length distribution of each strain at different temperatures**

| | **WT (22 x 30°C)** | ***gapR*/K59A (22 x 30°C)** | ***gapR*/R65A,K66A (22 x 30°C)** | ***gapR*/Q19R,L30P (22 x 30°C)** |
|---|---|---|---|---|
| mean length difference (µm) (IC - 95%) | 0.068 (0.018 to 0.118) | 0.085 (-0.023 to 0.193) | 0.736 (0.601 to 0.871) | 0.106 (-0.161 to 0.373) |

### REFERENCES


1. Ducret, A., Quardokus, E.M. and Brun, Y.V. (2016) MicrobeJ, a tool for high throughput bacterial cell detection and quantitative analysis. *Nat Microbiol*, **1**, 16077.
2. Rueden, C.T., Schindelin, J., Hiner, M.C., DeZonia, B.E., Walter, A.E., Arena, E.T. and Eliceiri, K.W. (2017) ImageJ2: ImageJ for the next generation of scientific image data. *BMC Bioinformatics*, **18**, 529.
3. Kanehisa, M. and Goto, S. (2000) KEGG: kyoto encyclopedia of genes and genomes. *Nucleic Acids Res*, **28**, 27-30.
4. Bertin, P., Benhabiles, N., Krin, E., Laurent-Winter, C., Tendeng, C., Turlin, E., Thomas, A., Danchin, A. and Brasseur, R. (1999) The structural and functional organization of H-NS-like proteins is evolutionarily conserved in gram-negative bacteria. *Mol Microbiol*, **31**, 319-329.
5. Buggy, J.J., Sganga, M.W. and Bauer, C.E. (1994) Characterization of a light-responding trans-activator responsible for differentially controlling reaction center and light-harvesting-I gene expression in *Rhodobacter capsulatus*. *J Bacteriol*, **176**, 6936-6943.
6. Madeira, F., Park, Y.M., Lee, J., Buso, N., Gur, T., Madhusoodanan, N., Basutkar, P., Tivey, A.R.N., Potter, S.C., Finn, R.D. *et al.* (2019) The EMBL-EBI search and sequence analysis tools APIs in 2019. *Nucleic Acids Res*, **47**, W636-W641.
7. Gao, Y., Foo, Y.H., Winardhi, R.S., Tang, Q., Yan, J. and Kenney, L.J. (2017) Charged residues in the H-NS linker drive DNA binding and gene silencing in single cells. *Proc Natl Acad Sci U S A*, **114**, 12560-12565.
8. Evinger, M. and Agabian, N. (1977) Envelope-associated nucleoid from *Caulobacter crescentus* stalked and swarmer cells. *J Bacteriol*, **132**, 294-301.
9. Guzman, L.M., Belin, D., Carson, M.J. and Beckwith, J. (1995) Tight regulation, modulation, and high-level expression by vectors containing the arabinose PBAD promoter. *J Bacteriol*, **177**, 4121-4130.
10. Ried, J.L. and Collmer, A. (1987) An nptI-sacB-sacR cartridge for constructing directed, unmarked mutations in gram-negative bacteria by marker exchange-eviction mutagenesis. *Gene*, **57**, 239-246.
11. Ricci, D.P., Melfi, M.D., Lasker, K., Dill, D.L., McAdams, H.H. and Shapiro, L. (2016) Cell cycle progression in *Caulobacter* requires a nucleoid-associated protein with high AT sequence recognition. *Proc Natl Acad Sci U S A*, **113**, E5952-E5961.
